# Inactivation of the DREAM complex mimics the molecular benefits of sleep

**DOI:** 10.1101/2024.06.26.600859

**Authors:** Isabela Santos Valentim, Francisco Javier Romero-Expósito, Katjana Schwab, Melike Bayar, Adiya Tauassarova, Konstantin Riege, Lisa Lange, Olivia Engmann, Martin Fischer, María Olmedo, Steve Hoffmann, Maria A. Ermolaeva

**Affiliations:** Leibniz Institute on Aging – Fritz Lipmann Institute (FLI), Beutenbergstrasse 11, 07745 Jena, Germany; Department of Genetics University of Sevilla, Av. Reina Mercedes s/n, 41012 Sevilla, Spain; Institute of Biochemistry and Biophysics, Center for Molecular Biomedicine (CMB), Friedrich Schiller University Jena, Hans-Knöll-Str. 2, Jena 07745, Germany; Cluster of Excellence Balance of the Microverse, Friedrich Schiller University Jena, Jena, Germany

## Abstract

Circadian clock disruption and lack of sleep impair organismal health, but remedies remain elusive. Here, we used multi-omics, molecular and functional assays in *C. elegans*, human retinal cells and mouse brain to identify the DREAM complex as a conserved mediator of sleep and clock benefits at the cellular level. We show that DREAM abundance is under circadian control with high levels seen during wakefulness. DREAM elevation confers increased chromatin compaction and shields DNA from damage while altering fundamental cellular processes such as translation, stress responses and OXPHOS. Conversely, DREAM levels are lowered during sleep enabling cellular maintenance and repair. When sleep or clock are altered, DREAM levels remain high, and repair activities remain suppressed triggering cellular and organismal deterioration that can be reversed by genetic and pharmacological inhibition of DREAM *in vivo* and *in vitro*. We thus reveal the potential of DREAM inhibitors to replicate the benefits of sleep in sleep-deprived and clock-impaired organisms.

## Introduction

Circadian clock impairment and sleep deprivation have disruptive impact on multiple aspects of organismal fitness from cognition to the immune response^1–3^. This multifaceted influence can partially be explained by the requirement of sleep for the fluid flow and molecular damage clearance in the brain^4^, with central nervous system (CNS) mediating a plethora of organismal responses^5^. An alternative hypothesis is that sleep deprivation and circadian disruption impact master regulatory pathways, which control fundamental aspects of the cellular physiology affecting multiple cell types. Identifying such core regulatory mechanisms could advance interventions to restore homeostasis during circadian misalignment and sleep deprivation. Here, we used RNAi against the *C. elegans PER* orthologue *lin-42* to induce persistent but moderate clock disruption as a mimetic of lifestyle-inflicted clock irregularities. We next applied this intervention in *C. elegans* models of proteostasis stress and accelerated aging followed by proteomics, molecular and functional assays. In parallel, we conducted transcriptomic analyses in mice with natural and altered sleep cycles, and in *PER1* knock-down human cells. These tests revealed the worm DRM complex (DP, Rb and MuvB) and its related mammalian DREAM complex (DP, RB-like, E2F4 and MuvB) as putative core regulators of cellular and molecular effects of clock and sleep. By temporal proteomics and transcriptomics tests we found DRM/DREAM abundance to be increased during natural wakefulness and in clock/sleep disrupted animals. Higher DREAM activity led to an increased abundance of histones implicated in chromatin condensation, and shielded DNA from damage simultaneously altering cellular maintenance and repair. Conversely, during sleep DREAM levels were lowered facilitating homeostasis and repair. When sleep and clock were disrupted, DREAM levels and histone abundance remained high leading to inferior repair and cellular decline that could be reversed by DREAM inhibitors. Our data suggests that DREAM is employed by the circadian cycle to condense the chromatin during wakefulness thus shielding DNA from the excessive damage, while during sleep low DREAM levels promote chromatin relaxation thus boosting repair activities including DNA repair. The circadian changes of DREAM abundance thus allow balancing DNA shielding with repair efficacy at the most optimal times of the circadian cycle, and we propose, based on functional evidence, that the restorative benefits of sleep can be mimicked by DREAM inhibitors provided in a timely manner.

## Main

### Moderate distortion of the circadian clock leads to tissue dysfunction in combination with cellular stress

In *C. elegans,* we employed RNAi-mediated gene inactivation to knock down a core component of the circadian clock *PER*/*lin-42*^6–10^. The complete loss of *lin-42* function was previously found to cause accelerated aging and early death in the nematodes^11^. This finding aligns with recent reports of reduced lifespan and increased cellular stress in sleep-deprived *D. melanogaster* and mice ^12^. Unlike complete clock inactivation, we focused on the organismal effects of partial clock impairment, akin to occasional loss of circadian function due to lack of sleep in mammals. We therefore used the RNAi-mediated gene knockdown (KD) approach instead of loss-of-function mutants. We found that *lin-42* RNAi treatment from the L1 larval stage disrupted the clock-dependent molting process of *C. elegans* (Figure 1A, B and S1A) without causing a reduction of lifespan (Figure 1C), consistent with moderate clock impairment. Because previous studies in mice and *D. melanogaster* suggested that sleep deprivation induces cellular stress^12^, we next asked if moderate circadian clock disruption would impair organ function in combination with additional stressors. Here, we used transgenic expression of aggregation-prone proteins to induce cellular stress in the body wall muscle of *C. elegans*. Specifically, we used nematodes expressing human amyloid-β peptide (*unc-54p::human Aβ_1-42_*) and polyglutamine coupled YFP protein (*unc-54p::Q40::YFP*) in body wall muscle^13,14^. Interestingly, *lin-42* inactivation from the L1 stage significantly worsened muscle function in both transgenic models, as seen by reduced motility (Figure 1D and S2A). A similar trend was seen in transgenic animals treated with *lin-42* RNAi from the pre-adult L4 stage (Figures S2B, C), demonstrating independence from developmental heterochronicity and showing that clock disruption during adulthood is sufficient to impair tissue function under stress^3,15,16^. At the same time, developmental clock disruption had a greater impact on lifelong tissue function, reflecting the critical role of sleep and coordinated quiescence during growth and development ^17–22^. Finally, RNAi knockdown of the downstream clock gene *lin-14*^23–25^ similarly reduced motility in *unc-54p::Q40::YFP* animals, like knockdown of the upstream regulator *lin-42* (Figure S2D), confirming clock disruption as the link between *lin-42* inactivation and stress-induced tissue failure. Collectively, we found that moderate but persistent clock impairment triggers organ dysfunction in combination with cellular stressors.

**Fig. 1.**
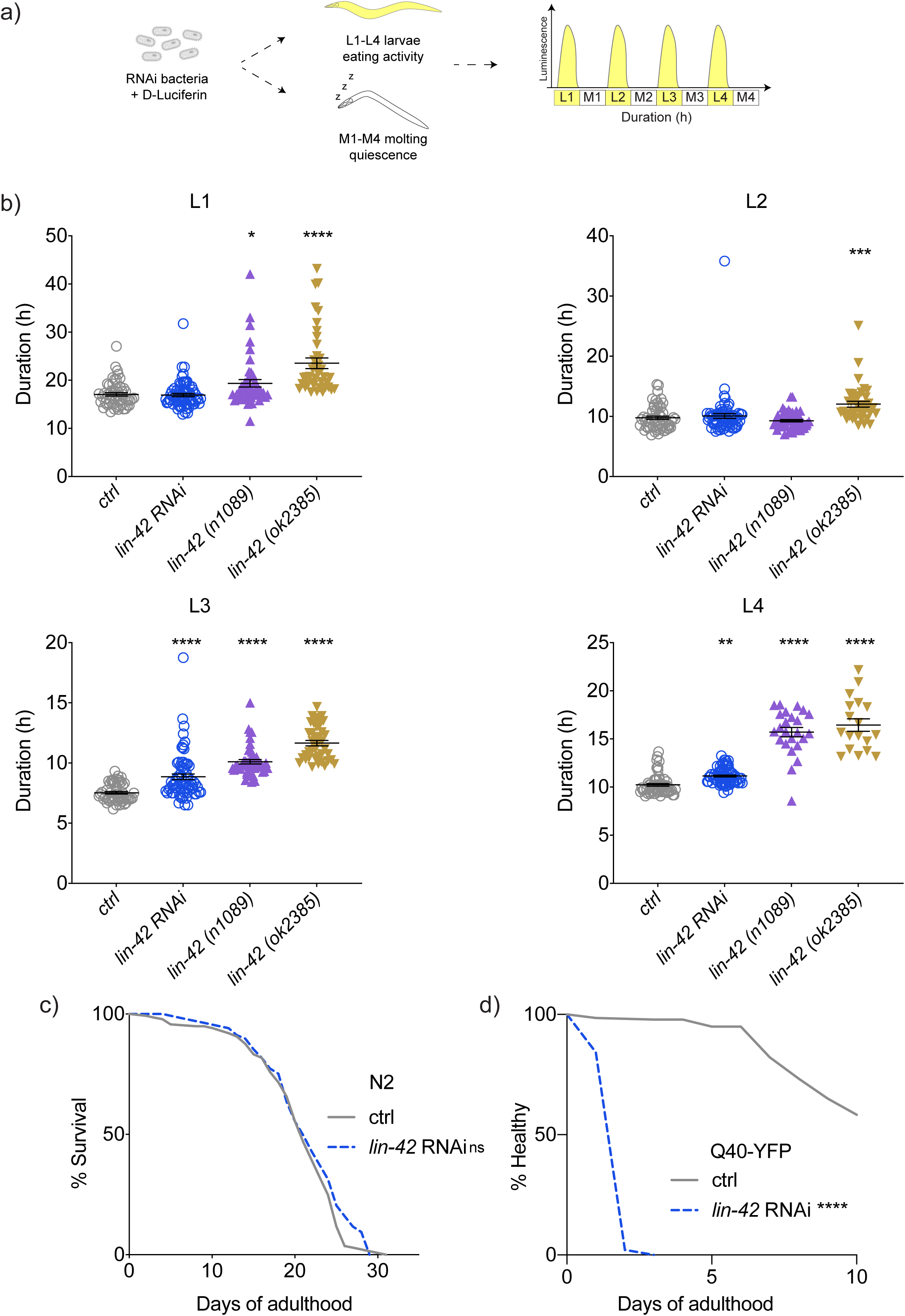
Knockdown of *lin-42*/PER impairs clock-controlled developmental timing and exacerbates aggregation-driven tissue failure. **(a)** Luminometry experimental design: Animals expressing a luciferase reporter of food intake were used. Feeding during larval stages triggers an increase of luminescence while quiescence and lack of feeding during molting stages lead to a decrease of signal. **(b)** Developmental timing (duration of the larval stages L1-L4) of control (ctrl), *lin-42* RNAi knock-down, *lin-42(n1089)* mutant and *lin-42(ok2385)* mutant animals is shown. Control (EV RNAi) and *lin-42* RNAi cohorts were established in the wild type (N2 Bristol strain) background. *lin-42(n1089)* mutant and *lin-42(ok2385)* mutant animals were fed with EV RNAi, and all worms were fed with D-Luciferin. Larval stage durations were quantified at 20°C based on relevant luminescence signals. Each dot represents one animal; average and S.E.M values are plotted. n=5-20 worms per condition in each trial, and results of 4 independent trials are combined. One-way ANOVA was used for the statistical assessment, two-tailed *p* values were computed, *-*p*<0.05; **-*p*<0.01; ***-*p*<0.001; ****-*p*<0.0001. **(c)** Wild-type (N2) worms were age-synchronized and fed with EV (ctrl) or *lin-42* RNAi from the L1 stage. Survival was scored daily. Animals were transferred to new plates every second day until adulthood day 10 (AD10), and every fourth day thereafter. n=140 worms per condition, graph is representative of three independent experiments. Significance was measured by the Log-rank Mantel-Cox test, two-tailed *p* values were computed. ns, not significant. **(d)** Age-synchronized populations of *unc-54*p::Q40::YFP animals were fed with EV (ctrl) or *lin-42* RNAi from the L1 stage. Paralysis was scored daily. Worms were transferred to new plates every second day. n=140 worms per condition, graph is representative of three independent experiments. Significance was measured by the Log-rank Mantel-Cox test, two-tailed *p* values were computed. ****-*p*<0.000001. Exact *n* numbers, *p*-values, and relevant statistical calculations for this and other figures are provided in the Statistics Source Data file.

### Clock disruption elicits expression changes of chromatin components and regulators of DNA metabolism

To elucidate the molecular mechanisms linking clock disruption to organ frailty, we performed unbiased proteomics analysis of the animals exposed to *lin-42* RNAi from L1 and L4 stages (Figure 2A and S4A) identifying 5,358 protein groups (Tables S1-S2). Principal component analysis (PCA) of whole proteomes revealed clear separation between *lin-42* RNAi-treated and control (EV) animals when treatment began at L1 (Figure 2B). A subsequent gene set enrichment analysis of the proteins differentially expressed between the two experimental groups revealed a significant enrichment of the terms associated with chromatin organization and DNA metabolism (Figure 2C and Table S3). In parallel, the unbiased volcano plot analysis performed on the same set of proteins revealed several histone variants as the most clearly up-regulated protein sub-group (Figure 2D). Together, these two independent and unbiased whole-proteome analyses suggest that chromatin changes may underlie the cellular effects of *lin-42* knockdown (Figure 2F). To identify the core regulators linking *lin-42* inactivation to chromatin re-organization, we next computationally extracted the top protein contributors to the PC1 separation between *lin-42* RNAi and EV control proteomes (Figure 2B, Table S4), and analyzed these by using the STRING protein-protein interaction assessment tool^26^. Interestingly, the strongest observed network included the key DRM complex component *lin-35*^27^, as well as known DRM/DREAM targets and associated proteins such as histones *his-41*/H2BC and *htz-1*/H2A.Z2, and histone binding proteins *hcp-6*/NCAPD3 and *capg-2*/NCAPG2^28–30^ (Figure S3, Table S5). Notably, DRM and its mammalian counterpart DREAM participate in chromatin remodeling and gene expression regulation^31,32^, in matching with the strong impact of clock disruption on chromatin components observed in Figures 2C and 2D. In addition to *lin-35* being a key contributor to proteome segregation along the PC1, most of individual DRM subunits and some of DRM-associated histones were upregulated in animals exposed to *lin-42* RNAi from both L1 and L4 stages (Figure 2E, Tables S6-S7). Moreover, comparable DRM/DREAM-associated proteins contributed substantially to the PC1-separation in both L1- and L4-treated cohorts (Figure S3 and S4A-C, Tables S4-S5). Overall, L1-initiated clock disruption caused stronger molecular changes, consistent with its greater impact on organismal health (Figure 1D, Figures S2A-C). These findings uncover the DREAM complex as a key molecular target of the circadian clock disruption, and demonstrate that the molecular effects of clock impairment are similar between development and adulthood, although they differ in magnitude.

**Fig. 2.**
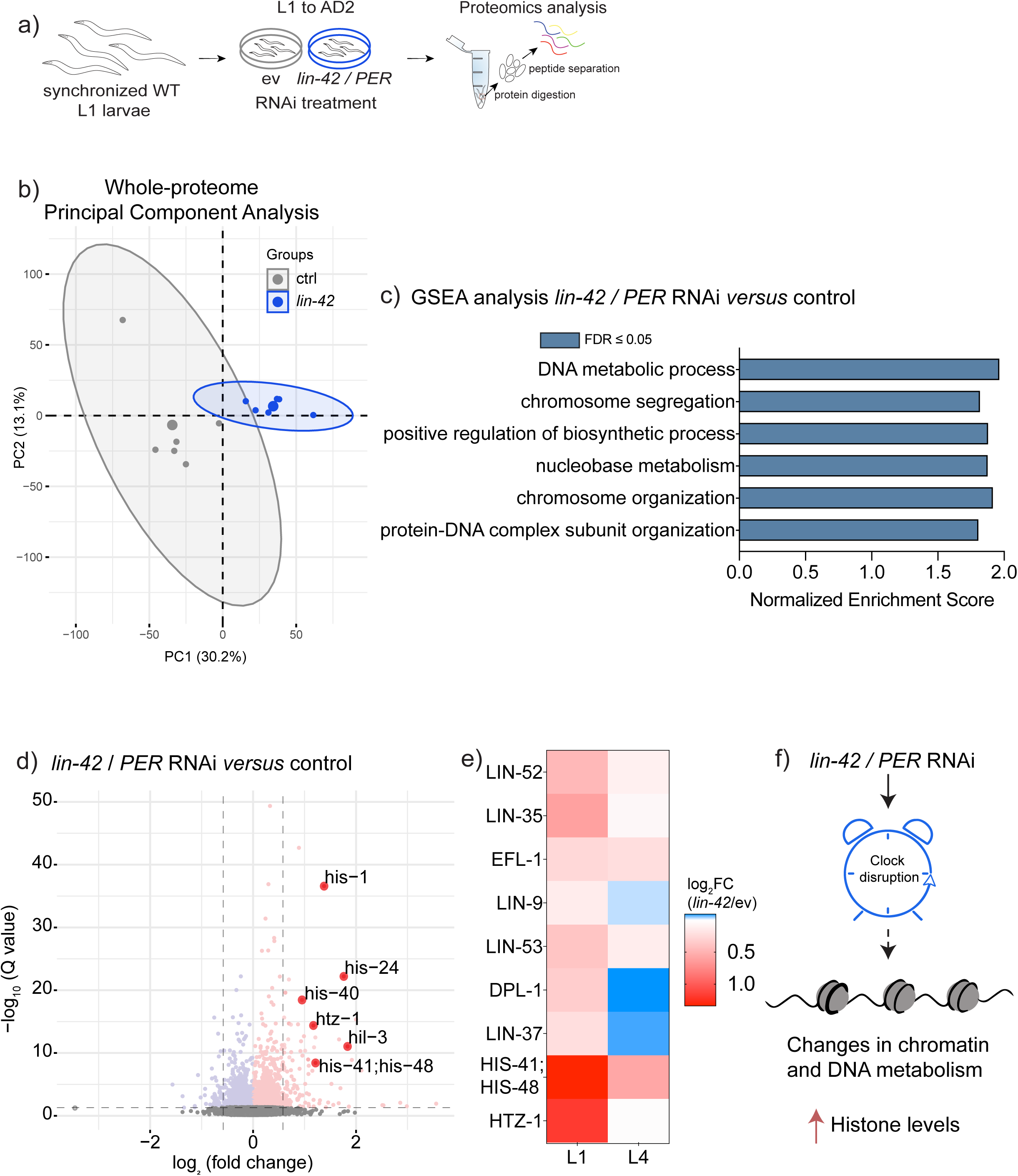
Clock disruption triggers changes in histone expression, chromatin organization, and DRM/DREAM complex abundance. **(a)** Experimental design. **(b)** Principal component analysis comparing whole proteomes of wild-type *C. elegans* treated with EV or *lin-42* RNAi from the L1 stage is shown. Proteins were extracted on AD2, and small dots are representative of independent replica samples, n=800 worms per replica. **(c)** Bar plot of significantly enriched (FDR ≤ 0.05) GO terms among proteins upregulated (Average Log_2_ expression FC>0) between *lin-42* and EV RNAi-treated animals is shown. **(d)** Volcano plot depicting differently expressed proteins between *lin-42* and EV RNAi-treated cohorts is shown, with histones highlighted in red. The horizontal dashed line indicates a Q-value cutoff of 0.05 and the vertical lines - a log_2_ expression fold change cut-off of ≤0.58. **(e)** Heatmap of selected DRM/DREAM subunits and interactors is shown. The color code refers to log_2_ expression fold change values between *lin-42* and EV RNAi-treated cohorts; RNAi treatments from both L1 and L4 stages are included. **(f)** Mechanistic model is presented. The complete data and calculations for panels **b**-**e** can be found in Tables S1, S3 and S6-7 respectively.

### DRM/DREAM complex subunits and histones mediate tissue dysfunction in response to circadian clock disruption

To validate the role of DRM/DREAM in the tissue dysfunction caused by circadian clock disruption, we performed co-knock downs of *lin-42* and individual DRM subunits by double RNAi exposure of the *unc-54p::Q40::YFP* animals. In these tests, we focused on RNAi treatment initiated at the L1 larval stage because of the stronger functional response observed in this setting previously (Figure 3A). Strikingly, we found that RNAi-mediated depletion of DRM subunits *lin-9*, *lin-54* and *lin-53* reversed the motility defects caused by the *lin-42* gene knock down in the *unc-54p::Q40::YFP* animals, with *lin-53* inactivation showing the strongest effect (Figure 3B-D). Notably, our double RNAi approach was effective in suppressing both genes simultaneously as seen with the example of *lin-42*/*lin-53* co-suppression (Figure S5A). At the same time, knock down of other canonical DRM subunits such as *lin-37*, *lin-52* and *lin-35* did not affect tissue dysfunction elicited by the *lin-42* RNAi (Figure S5B-D) demonstrating selective requirement of the distinct DRM/DREAM subunits in the effects of circadian clock. Moreover, we showed that RNAi-mediated depletion of the *htz-1*/*H2A.Z2* histone variant found to be associated with the DRM/DREAM complex in our STRING analysis (Figure S3) and in literature^29^, also rescued enhanced paralysis of the *unc-54p::Q40::YFP* nematodes treated with the *lin-42* RNAi (Figure 3E). Collectively, our results indicated that the enhanced activity of specific DRM/DREAM complex variants might mediate the negative impact of clock inactivation on tissue homeostasis under stress. Our data also suggested that increased abundance of DRM/DREAM-associated histones might play a role in this process by broadly altering chromatin composition and gene expression with possible effect on the adaptive cellular activities. In line with this model of non-selective gene repression, YFP expression was progressively reduced with age in *unc-54p::Q40::YFP* animals exposed to *lin-42* RNAi (Figure S6A and B), and this effect was reversed in worms co-treated with the anti-DRM *lin-53* RNAi. At the same time, relative YFP aggregation was progressively increased with age in the *lin-42* RNAi exposed animals, and this distortion was also reversed by the co-knock down of the *lin-53* DRM subunit in line with respectively impaired and restored ability to counteract proteostasis stress (Figure 3F).

**Fig. 3.**
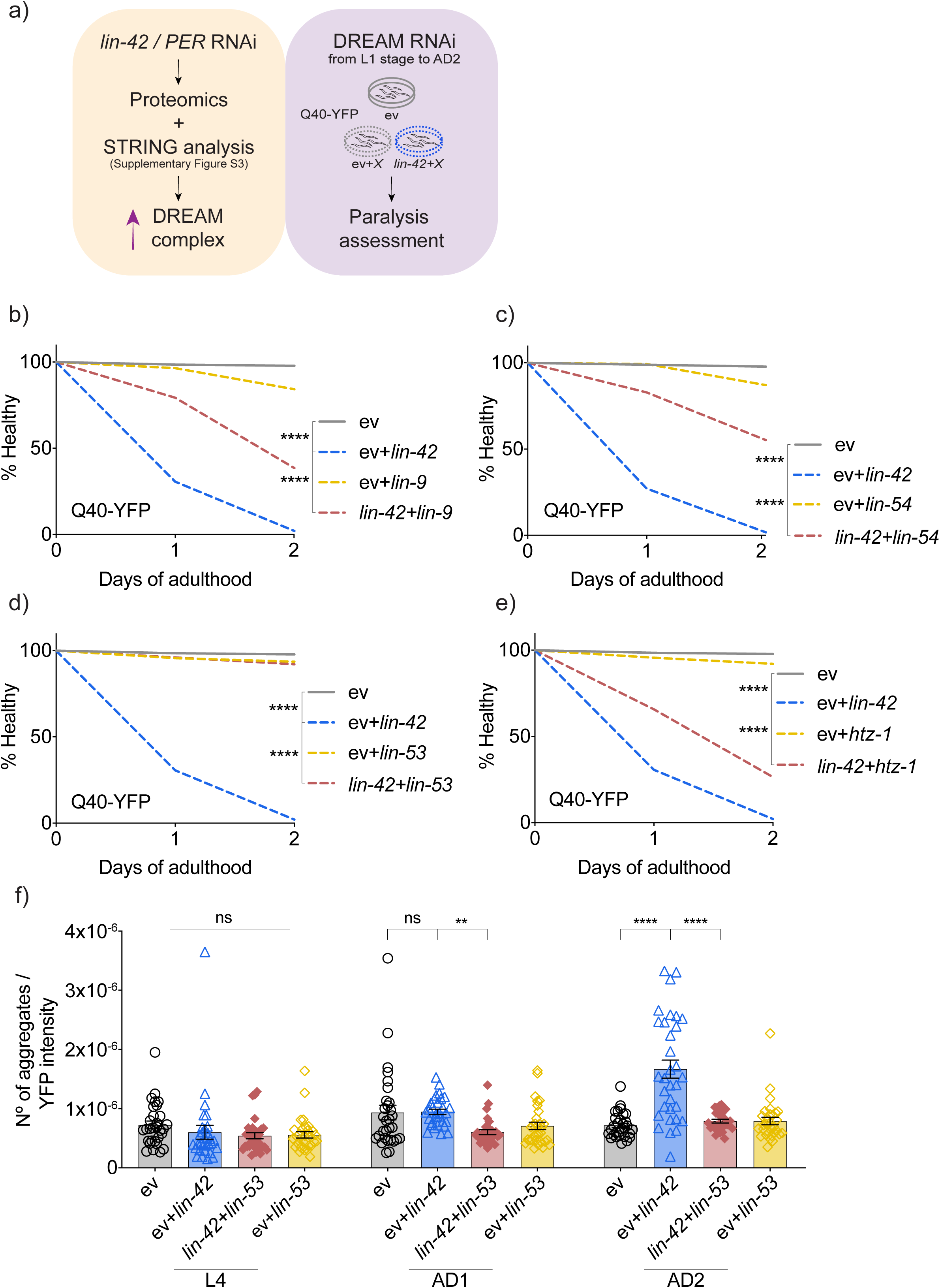
Inactivation of the DRM/DREAM complex reverses functional impairments induced by clock disruption. **(a)** Left caption reminds of elevated DRM/DREAM abundance in clock-impaired animals; right caption depicts the experimental design. **(b-e)** Age-synchronized *unc-54*p::Q40::YFP animals were fed with EV or *lin-42* RNAi in combination with RNAis targeting specific subunits/interactors of the DRM/DREAM complex from the L1 stage, paralysis was scored daily. Co-targeting with *lin-9* **(b)**, *lin-54* **(c)**, *lin-53* **(d),** and DRM/DREAM interactor *htz-1* **(e)** is shown. n=140 worms per condition, graphs are representative of at least three independent experiments. Significance was measured by the Log-rank Mantel-Cox test, two-tailed *p* values were computed. ****-*p*<0.0001. **(f)** *unc-54*p::Q40::YFP worms were age-synchronized, fed with indicated RNAi constructs from the L1 stage, and imaged at three different time points: L4, adulthood day 1 (AD1), and AD2. The number of aggregates divided by respective YFP intensity is shown, n=26-32 worms per condition, graph is representative of three independent experiments. Error bars are S.E.M. Significance was measured by one-way ANOVA within each age group separately and applying Sidak’s multiple comparison test, two-tailed *p* values were computed. **-*p*<0.01; ****-*p*<0.0001; ns, not significant.

### Clock disruption interferes with fundamental cellular activities in a DRM/DREAM-dependent manner

We next used Western blot analysis to test if the increased abundance of histones in *lin-42* KD animals was indeed regulated by DRM/DREAM. Consistent with the proteomics data shown in Figure 2D, histone abundance was elevated following *lin-42* RNAi exposure, and this elevation was reversed by the additional knock down of *lin-53* in all cases (Figure 4A and B, Figures S7 and S8A). Notably, the strongest difference in protein abundance between control, *lin-42* KD and *lin-42/lin-53* co-knockdown cohorts was seen for H1 and H2B histone variants, known to regulate and enhance chromatin compaction (Figure 4A and B, Figure S7A-B)^33,34^, and their differential expression was observed also at the mRNA level (Figure S8B). These results indicate that DREAM/DRM facilitates higher histone levels (Figure 4L), likely leading to enhanced chromatin compaction.

**Fig. 4.**
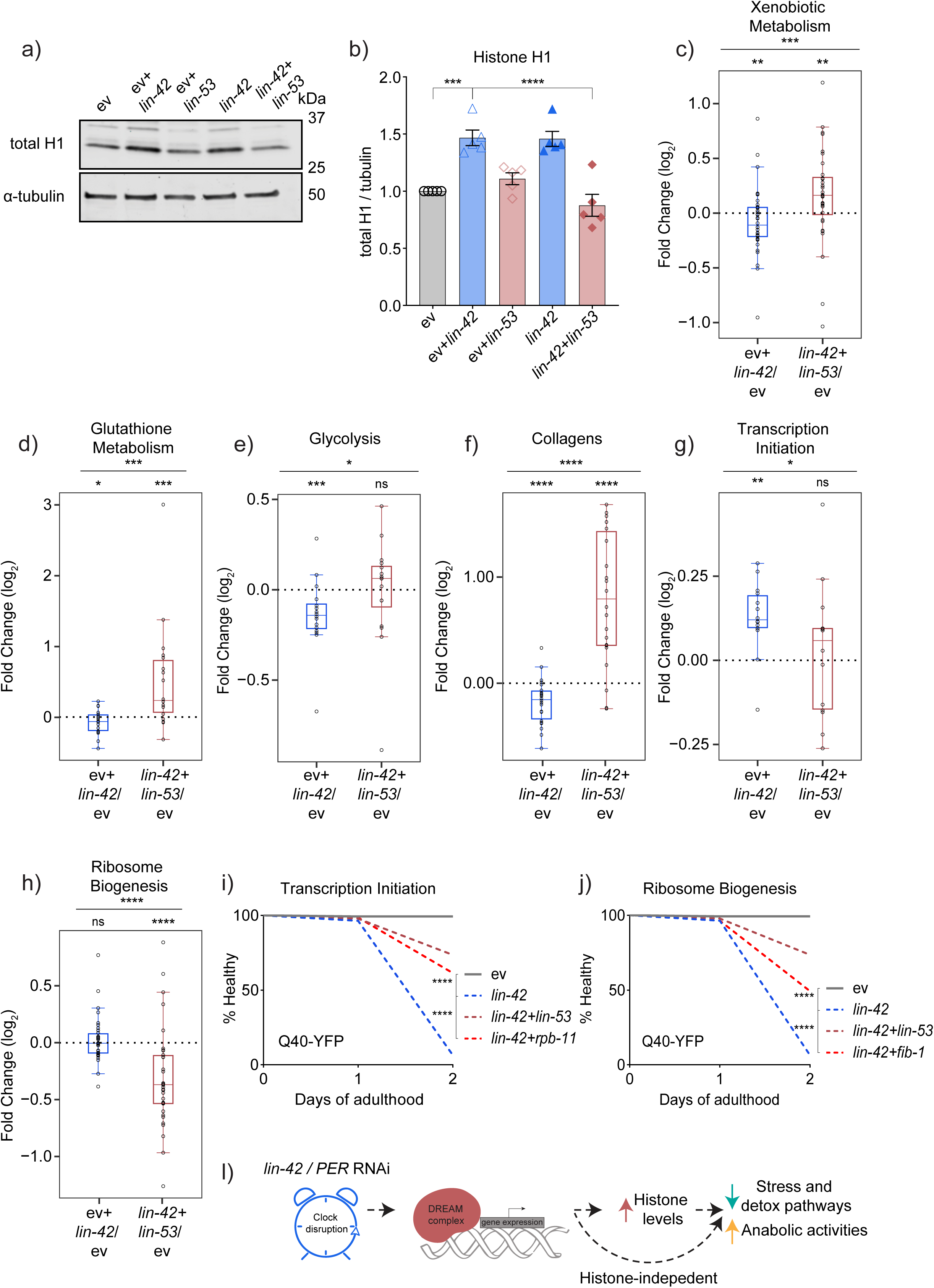
Clock disruption interferes with fundamental cellular activities in a DRM/DREAM-dependent manner. **(a)** Wild-type worms were age-synchronized and treated with indicated RNAi combinations from the L1 stage and until AD2. Expression of histone H1 was measured by immunoblot using tubulin as a loading control, full gel images are shown in Figure S7. **(b)** Quantification of 5 independent experiments represented by (**a**) is shown; n=800 worms in each condition. Error bars are S.E.M. Significance was measured by one-way ANOVA, two-tailed *p* values were computed. ***-*p*<0.001, ****-*p*<0.0001, ns, not significant. **(c-h)** Worms were treated with indicated RNAi combinations and control (EV) RNAi from the L1 stage until AD2. Protein expression was resolved by proteomics. Box plots showing relative expression (*EV+lin-42* over *EV,* and *lin-42+lin-53* over *EV* comparisons) of proteins belonging to xenobiotic metabolism **(c)**, glutathione metabolism **(d)**, glycolysis **(e)**, collagens **(f)**, transcription initiation **(g)** and ribosome biogenesis **(h)** are presented. Individual proteins are shown as dots, the median fold change of each group is shown as a horizontal line; the upper and lower limits of the box plot indicate the first and third quartile and the whiskers extend 1.5 times the interquartile range from the limits of each box. n=800 worms per condition and 4 independent populations were measured in each case. Within each box plot, the significance was assessed by the Mann-Whitney Wilcoxon rank-sum test, and Wilcoxon test was used for the comparison between the two sets of fold changes. Two-tailed *p* values were computed in all cases. *-*p*<0.05; **-*p*<0.01; ***-*p*<0.001; ****-*p*<0.0001. The lists and data of all included proteins can be found in Tables S8 (complete data) and S9 (box plot data). **(i-j)** Age-synchronized *unc-54*p::Q40::YFP animals were fed with EV or *lin-42* RNAi in combination with RNAis targeting specific genes implicated in transcription initiation and ribosome biogenesis from the L1 stage, paralysis was scored daily. Co-targeting with *rpb-11* **(i)** and *fib-1* **(j)** is shown. n=140 worms per condition, graphs are representative of three independent experiments. Significance was measured by the Log-rank Mantel-Cox test, two-tailed *p* values were computed. ****-*p*<0.0001. **(l)** Mechanistic model is depicted.

Subsequently, we asked what molecular pathways become de-regulated in *lin-42* knockdown nematodes and re-instated in *lin-42/lin-53* double knockdown animals, representing molecular targets affected by the circadian clock disruption in a DRM/DREAM-dependent manner. *lin-53* inactivation was chosen due to its strongest ability of reversing the negative health effects of the *lin-42* KD (Figure 3D), and because *lin-53*/*RBBP4* was reported to mediate the interaction between DRM/DREAM and histones^35,36^. The comparative proteomics analysis of *lin-42* KD, *lin-42/lin-53* co-knockdown and EV RNAi control-exposed cohorts (Figure S9A and Table S8) strikingly revealed simultaneous alterations of multiple mechanisms implicated in both basal homeostasis of the cell and in adaptive stress responses. For instance, detoxification mechanisms such as xenobiotic metabolism (Figure 4C and Table S9) and pathways implicated in the antioxidant response (Figures 4D, S9B-D and Table S9) were found to be suppressed by *lin-42* KD and restored by the co-knockdown of *lin-53*. The antioxidant response was represented by simultaneous de-regulation of four distinct metabolic entities implicated in counteracting oxidative stress: aldehyde dehydrogenase protein family (Figure S9B)^37^, glutathione metabolism (Figure 4D)^38^, tryptophan catabolism by kynurenine pathway (Figure S9C) with intermediates known to exert ROS scavenging roles^39–41^, and short-chain dehydrogenase protein family implicated in xenobiotic and antioxidant defense responses (Figure S9D)^42,43^. The outstanding enrichment of antioxidant mechanisms among the DRM-dependent clock targets fits well with oxidative stress being the primary cause of the early demise in sleep deprived *Drosophila* and mice^12^. At the same time, xenobiotic detoxification responses were previously found to be de-regulated in the livers of *Per1*/*Per2* double-knockout mice suffering from clock impairment^44^, in line with our present findings. In addition, metabolic plasticity pathways such as glycolysis and fatty acid β-oxidation^45^ were diminished by *lin-42* KD and reinstated by *lin-53* co-inactivation (Figure 4E and S9E, Table S9), while key bioenergetics activities of mitochondria – TCA cycle and OXPHOS, were dampened by *lin-42* inactivation and partially rescued by *lin-53* RNAi co-treatment (Figure S9F and G, Table S9). Finally, the expression of ECM collagens was downregulated in *lin-42* RNAi-exposed animals and restored by *lin-42*/*lin-53* co-knockdown (Figure 4F, Table S9). Notably, both decline of metabolic plasticity and reduced expression of collagens are associated with loss of organ function during aging^45–47^, establishing a link between clock disruption and accelerated tissue impairment. Moreover, an impairment of metabolic adaptive capacity was previously observed in *Per1*/*Per2* deficient mouse livers^44^. The cell-protective interplay between metabolic and detoxification activities regulated by DRM/DREAM and clock can also be envisaged because tryptophan catabolism by kynurenine pathway is one of the key cellular sources of nicotinamide adenine dinucleotide (NAD+)^48^– an energy carrier and enzymatic co-factor with key roles in mitochondrial maintenance and bioenergetics fitness, and with capacity to delay metabolic aging^48^. The strong enrichment of metabolic pathways among DRM-dependent clock targets is in line with the large body of human clinical data designating clock and sleep dysfunction as confounding factors of metabolic diseases such as diabetes, obesity and cardio-vascular disorders^49,50^.

Interestingly, basal cellular activities such as Pol II transcription and splicing were differentially affected in our experimental system showing upregulation following clock distortion and re-blunting by co-knock down of *lin-53* (Figure 4G and S9H, Table S9). Of note, increased speed of Pol II transcription and elevated splicing were recently revealed to be hallmarks and functional drivers of physiological aging^51^, and the interference of the *lin-42* KD with both processes provided yet another putative *lin-53*/DRM-dependent link between clock disruption and premature organismal demise. Finally, we observed mitochondrial ribosome, cytosolic ribosome, ribosome biogenesis and translational proteins to be strongly downregulated by *lin-42/lin-53* co-knock down (Figure 4H and S9I-L, Table S9) suggesting that DRM/DREAM modulates these essential activities in the context of clock disruption. Because inhibition of translation is known to improve proteostasis by reducing protein folding stress^52–54^, the strong blunting of this process at multiple levels likely contributes to superior capacity of *lin-42*/*lin-53* double KD to reduce polyQ aggregates compared to *lin-42* only inactivated worms (Figure 3F).

To determine whether the mechanisms identified by proteomics played a functional role in the impairment of motility by clock disruption, we focused on the pathways upregulated by clock breakdown in a DREAM-dependent manner, e.g. transcriptional initiation, ribosome biogenesis, and translation. We suppressed these pathways using specific RNAi treatment in clock-disrupted *unc-54p::Q40::YFP* transgenic animals. RNAis against RNA polymerase II subunit *rpb-11* and TFIIH core complex helicase subunit *xpd-1* were used to attenuate transcription, while initiation factors *ife-2* and *ife-3* were targeted to suppress translational initiation, and RNA polymerase I subunit *rpoa-2* and pre-rRNA processing factor fibrillarin *fib-1* were targeted to interfere with ribosome biogenesis. Strikingly, all RNAi treatments mimicked the restorative effect of DREAM inhibition in the context of clock impairment (Figures 4I, J and S10), showing that DREAM-dependent pathway alterations are functional drivers of the negative changes caused by clock disruption. Notably, the pathways highlighted in Figures 4 and S9 are not among the canonical DRM/DREAM targets, which were not covered in our analysis. Moreover, by comparing the list of clock and DREAM co-targets depicted in Figures 4 and S9 to the previously published DREAM ChiP-Seq *C. elegans* dataset^29^, we determined that only few of these genes are located in the close proximity to the relevant genomic DREAM binding sites (Table S10). This finding speaks in favor of indirect chromatin- and histone-dependent regulation of these targets by DRM/DREAM, which is also supported by all of the functional and molecular data we obtained thus far. Overall, we found that clock disruption causes multiple alterations of basal and adaptive cellular functions that are reduced by co-knockdown of DRM/DREAM likely by affecting histone levels and chromatin structure (Figure 4L). The amplitude of changes in the individual adaptive pathways was not drastic, but in combination, these alterations have the potential to impede cellular homeostasis and flexibility at multiple levels contributing to health decline associated with clock and sleep impairments.

### DREAM expression fluctuates between sleep and wakefulness in mammals, impacting fundamental homeostatic mechanisms

We next asked if DREAM expression changes during the natural sleep-wake cycle by analyzing brain (frontal cortex) transcriptomes of young mice^55^. Interestingly, we found DREAM expression to be lowered during sleep and elevated during wakefulness (Figures 5A and S11B, 11D (right panel)), while sleep deprivation altered the circadian DREAM dynamics (Figures 5B and S11D, (left panel)) and prevented distinct DREAM subunits from being timely downregulated (Figure 5B), similar to observations in clock-impaired *C. elegans*. To validate if DREAM expression is under active circadian control, we performed qPCR on cDNA isolated from brain tissue (hippocampi) of mice following exposure to the sleep hormone melatonin that increases the amplitude of circadian responses^56^. We found expression of all tested DREAM subunits to be boosted by melatonin during the awake phase of the circadian cycle (Figures 5C and S12), suggesting that daily fluctuations of DREAM expression are part of the circadian response.

**Fig. 5.**
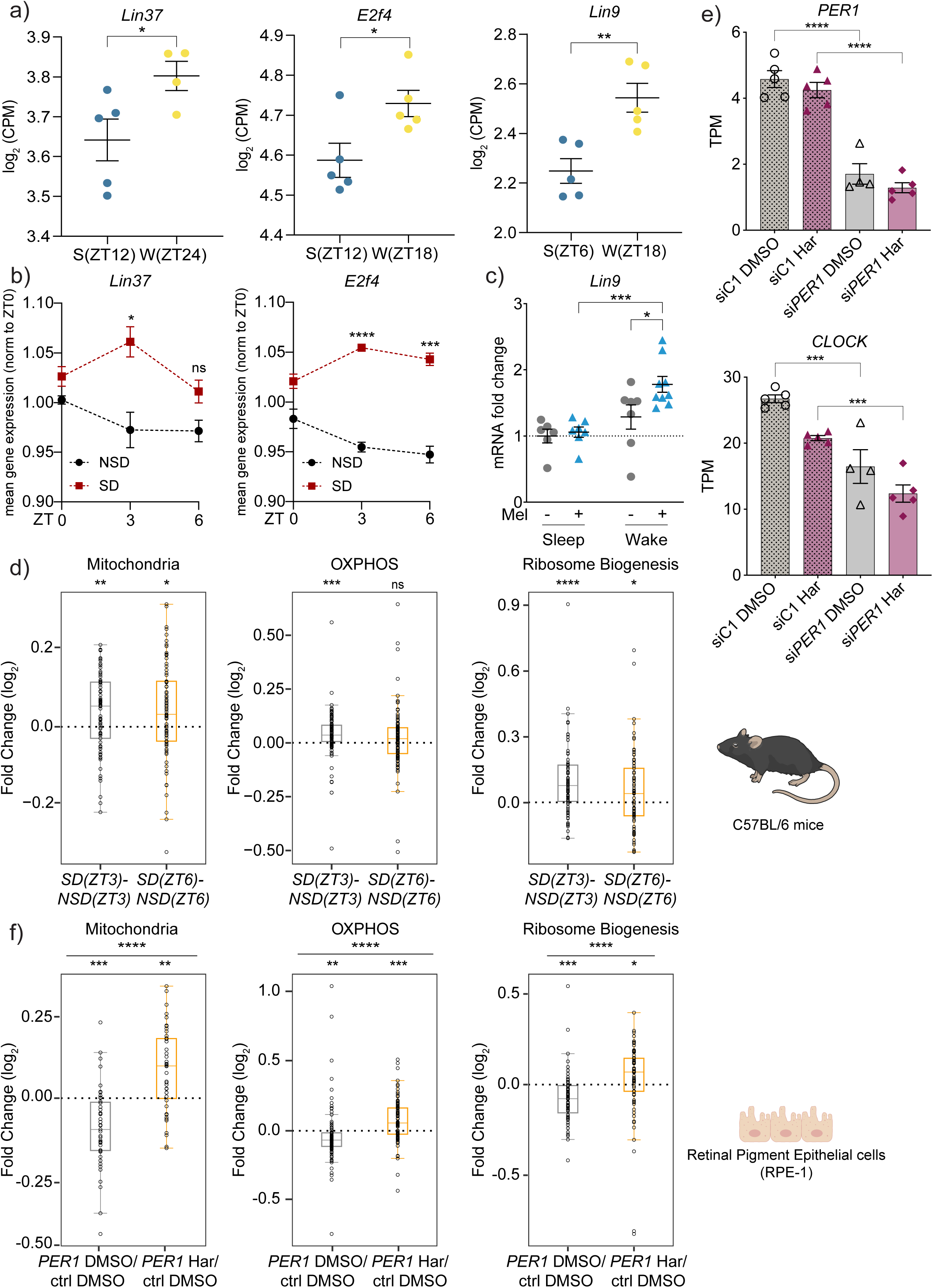
Circadian fluctuation of DREAM expression impacts fundamental homeostatic mechanisms in mammals. **(a-b)** Transcriptomic analysis of mouse cortex samples, collected during sleep, wakefulness and sleep deprivation (Fig. S11), and focusing on DREAM subunit expression. **(a)** Expression of *Lin37*, *E2f4* and *Lin9* subunits during sleep (blue circles) and wakefulness (yellow circles) is shown as Log_2_ CPM, each circle represents an independent sample. n= 3-5 in each condition, mean and S.E.M values are presented. Significance was measured by unpaired t-test at indicated ZTs. *-*p*<0.05; **-*p*<0.01. **(b)** The dynamics of *Lin37* and *E2f4* expression during early sleep (NSD) and corresponding sleep deprivation time points (SD) is shown. The Log_2_ CPM values at ZT 3 and 6 (NSD and SD) were normalized to corresponding ZT 0 values. n= 3-5 in each condition, mean and S.E.M values are presented. Significance was measured by unpaired t-test at indicated ZTs. *-*p*<0.05; **-*p*<0.01; ***-*p*<0.001; ****-*p*<0.0001; ns, not significant. The complete dataset and the normalization calculation can be found on Table S11 and S12. **(c)** Wild-type mice were treated with melatonin or vehicle for 2 weeks (experimental scheme FigS12), hippocampi were collected at both light and dark phases. Expression analysis of *Lin9* gene was performed by RT-qPCR. Data is shown as mRNA expression fold changes calculated by the ΔΔCt method. Significance was measured by multiple comparisons two-way ANOVA with a Bonferroni correction. n= 6-9 in each condition, mean and S.E.M values are presented. *-*p*<0.05; **-*p*<0.01; ***-*p*<0.001; ****-*p*<0.0001; ns, not significant. **(d)** Box plots showing relative expression (Log_2_ CPM SD (ZT3) - Log_2_ CPM NSD (ZT3) and Log_2_ CPM SD (ZT6) – Log_2_ CPM NSD (ZT6) comparisons) of genes belonging to Mitochondria, OXPHOS, and Ribosome Biogenesis. Individual genes are shown as dots, the median fold change of each group is shown as a horizontal line; the upper and lower limits of the box plot indicate the first and third quartile and the whiskers extend 1.5 times the interquartile range from the limits of each box. n=3-5 mice per condition were measured in each case. Within each box plot, the significance was assessed by the Mann-Whitney Wilcoxon rank-sum test, and the Wilcoxon test was used for the comparison between the two sets of fold changes. Two-tailed *p* values were computed in all cases. *-*p*<0.05; **-*p*<0.01; ***-*p*<0.001; ****-*p*<0.0001. The lists and data of all included genes can be found in Tables S13 (complete data) and S14 (box plot data). **(e)** Human retinal pigment epithelial cells (RPE) cells were treated with control siRNA (siC1, 10 nM) or siRNA against *PER1* (si*PER1*, 10 nM) for 48h and co-treated with Harmine (Har, 10μM) or DMSO (vehicle control) for 20h. Expression analysis of *PER1* and *CLOCK* genes by mRNA sequencing with graphs showing transcript per million values (TPM). Significance was measured by one-way ANOVA within each group separately, and applying Sidak’s multiple comparisons test, two-tailed *p* values were computed. *-*p*<0.05; **-*p*<0.01; ***-*p*<0.001; ****-*p*<0.0001; ns, not significant. n=4-5 independent samples. **(f)** Box plots showing relative expression (*PER1* DMSO versus ctrl DMSO and *PER1* Harmine versus ctrl DMSO comparisons) of genes belonging to Mitochondria, OXPHOS, and Ribosome Biogenesis are presented. Individual genes are shown as dots, the median fold change of each group is shown as a horizontal line; the upper and lower limits of the box plot indicate the first and third quartile and the whiskers extend 1.5 times the interquartile range from the limits of each box. Within each box plot, the significance was assessed by the Mann-Whitney Wilcoxon rank-sum test, and the Wilcoxon test was used for the comparison between the two sets of fold changes. Two-tailed *p* values were computed in all cases. *-*p*<0.05; **-*p*<0.01; ***-*p*<0.001; ****-*p*<0.0001. The lists and data of all included genes can be found in Tables S16 (complete data) and S17 (box plot data).

We next asked if impaired DREAM lowering under sleep deprivation impacts fundamental cellular activities in mice as it does in worms. Here, we compared brain transcriptomes of sleep deprived and sleep intact mice during the circadian window associated with naturally deeper sleep (Figure S11A). Of note, the same transcriptomic data and time window were used for the assessment of the relative DREAM dynamics between sleep and sleep deprivation in Figures 5 and S11. We found mitochondrial content, OXPHOS, glycolysis as well as ECM components (collagens), detoxification pathways, ribosome biogenesis and content, and splicing to be altered by sleep deprivation (Figures 5D and S13), similar to findings in clock-impaired *C. elegans*. Notably, recent studies uncovered that mitochondrial removal and the suppression of stress-inducing OXPHOS activity are central to the brain’s restorative processes during sleep^57,58^. Our findings reveal that both mitochondrial and OXPHOS gene expression failed to be dampened during sleep deprivation (Figure 5D), alongside alterations in DREAM abundance. Conversely, glycolysis became downregulated, mirroring results in *C. elegans* (Figure S13A). In addition, ribosome biogenesis was previously shown to be reduced during sleep^59^, likely alleviating energy demands and reducing protein folding pressure^60^. However, we observed that this homeostatic shift was impaired under sleep deprivation (Figure 5D and S13B). Moreover, detoxification activities such as tryptophan metabolism, glutathione metabolism and the xenobiotic detoxification pathway (Figure S13D-F) were dampened by sleep deprivation, similar to changes in clock impaired nematodes. Tryptophan turnover plays an additional role in regulating sleep quality in mammals^61^. Finally, de-regulation of brain collagens, as seen during sleep deprivation (Figure S13G), has been previously linked to neurodegeneration and dementia in humans^62^. We thus show that DREAM expression is high during wakefulness and low during natural sleep. We also demonstrate that DREAM expression fails to be timely lowered in sleep deprived mice similar to clock-impaired nematodes, and this failure correlates with de-regulation of similar homeostatic mechanisms as in clock-disrupted worms.

We next turned to a human cell culture system to test if *PER* inactivation is indeed sufficient to disrupt circadian clock and if inhibition of the DREAM complex can alleviate clock-related molecular impairments in the background of *PER* deficiency. Here we used siRNA-mediated knockdown of *PER1* in human retinal pigment epithelial (RPE) cells, and we used the DYRK1A inhibitor Harmine to pharmacologically suppress DREAM function, as seen in previous reports^63,64^. *PER1* was chosen as the key cycling *PER* paralogue across mammalian tissues as shown in a recent report^65^. We found that *PER1* knockdown was robust (Figures 5E and S14A) and indeed sufficient to cause clock impairments as seen by altered expression of *CLOCK* and *CRY2* core clock genes in qPCR and transcriptomic tests (Figures 5E and S14A). Importantly, Harmine did not restore the clock in *PER1*-depleted cells (Figures 5E and S14A), confirming our hypothesis that the restorative effect of DREAM inhibition on the homeostasis of clock-disrupted cells is independent of clock reinstatement and likely occurs downstream of an impaired clock.

Subsequently, we turned to deeper high-throughput transcriptomic analysis of *PER1*- and Harmine-treated RPE cells to identify the key homeostatic mechanisms regulated by clock in a DREAM-dependent manner in human cells (Table S16). Strikingly, our findings were similar to observations obtained in the nematode and mouse models. For instance, we found that mitochondrial and OXPHOS genes are downregulated by clock disruption and reinstated by co-inhibition of DREAM also in human cells (Figure 5F, Table S17). Moreover, ribosome and ribosome biogenesis genes, and genes encoding the components of the transcriptional machinery showed opposite expression dynamics between clock-disrupted and DREAM co-inhibited cells (Figures 5F and S14B-C, Tables S17-S18). These results matched the observations in nematodes and mice, and clearly suggested that similar fundamental cellular activities are altered by clock impairment in a DREAM-dependent fashion across species. Additionally, we found that co-administration of Harmine in *unc-54p::Q40::YFP* animals exposed to *lin-42* RNAi indeed reversed the paralysis induced by clock disruption in this model (Figure S14D and E). These results demonstrate that pharmacological inhibition of DREAM is a promising *in vivo* tool for rescuing the health of organisms affected by sleep and clock disruption.

### High DREAM abundance confers DNA shielding during wakefulness at the cost of inferior repair

We then explored the physiological significance of the circadian fluctuations of DREAM. Drawing from our findings that DREAM levels are elevated during wakefulness and contribute to increased histone abundance and likely histone loading, as well as literature evidence showing that histones and chromatin protect DNA from damage^66,67^ and that oxidative and DNA damage are heightened during wakefulness^57,68^, we hypothesized that the transiently high levels of DREAM serve to shield genomic DNA from the excessive damage associated with wakefulness. Indeed, inhibition of DREAM in cycling human cells led to elevated expression of *P21* (Figure 6A) and *P53* target genes^69^ (Figure 6B, Table S19) in line with the previous reports^69,70^ and indicative of genotoxic stress. At the same time, boosting of DREAM levels by *lin-42* RNAi exposure protected *C. elegans* from the larval arrest triggered by ultraviolet B treatment known to induce helix distorting DNA lesions^71^ (Figures 6C and S15), suggesting that DREAM elevation indeed protects DNA from the excessive damage. On the other hand, previous reports found that DREAM inhibition upregulates a plethora of DNA repair pathways^72,73^ and consistently, we found markers of genotoxic stress (*P21* and *Gadd45b*^74^) to be lowered during sleep in parallel with the reduction of DREAM expression, while representative DNA repair marker *Ddb2*^75^ was upregulated (Figure 6D-F, left panel). Conversely, sleep deprivation hindered the lowering of *P21* and *Gadd45b* expression levels (Figure 6D and E, right panel) like it impeded the lowering of DREAM expression, while *Ddb2* failed to be upregulated in this context (Figure 6F, right panel). Collectively, our findings suggest that DRM/DREAM has a dual role in protecting the genome integrity during the circadian cycle: during wakefulness high levels of DRM/DREAM likely shield DNA from damage by increasing histone abundance, while during sleep DREAM levels are lowered allowing DNA repair to take place along with other repair and maintenance activities that remain suppressed during wakefulness as a trade-off of DNA protection. In fact, previous reports indeed show that maintenance and repair mechanisms from autophagy to DNA repair are most effective during sleep^76–78^. Under sleep deprivation failure to downregulate DREAM likely interferes with repair, but as shown above we can mitigate this impairment with DREAM inhibitors such as Harmine.

**Fig. 6.**
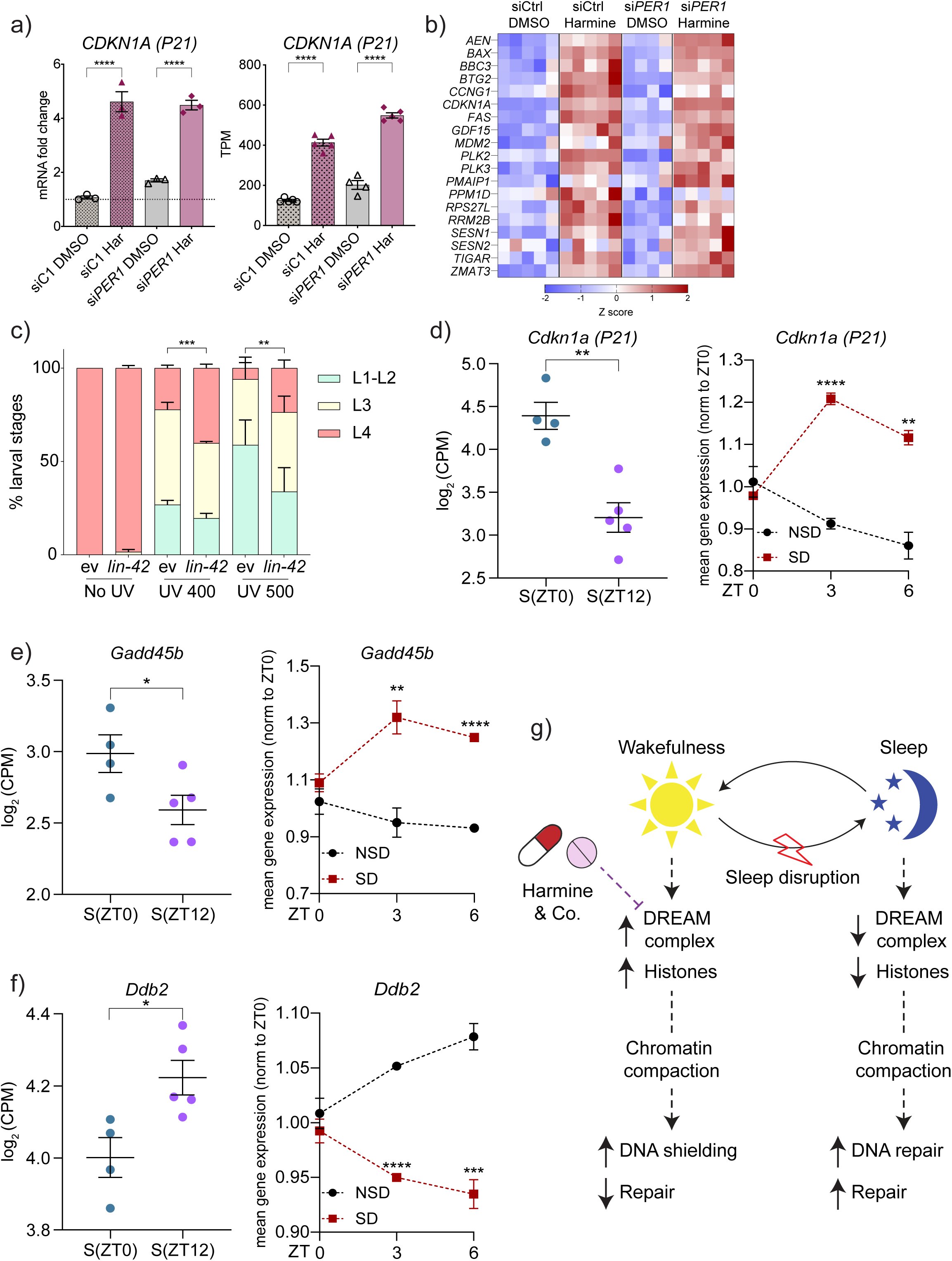
DREAM elevation confers DNA shielding during wakefulness at the cost of inferior repair. **(a)** Human retinal pigment epithelial cells (RPE) cells were treated with control siRNA (siC1, 10 nM) or siRNA against *PER1* (si*PER1*, 10 nM) for 48h and co-treated with Harmine (Har, 10μM) or DMSO (vehicle control) for 20h. **(a)** Expression analysis of *CDKN1A* gene was performed by RT-qPCR (left panel) and mRNA sequencing (right panel) in the same cell extracts. Data is shown as mRNA expression fold changes calculated by the ΔΔCt method and transcript per million values (TPM) respectively. Data in the left panel is representative of 5 independent trials, and 4-5 independent samples were combined in the right panel. Significance was measured by one-way ANOVA within each group separately, and applying Sidak’s multiple comparisons test, two-tailed *p* values were computed. *-*p*<0.05; **-*p*<0.01; ***-*p*<0.001; ****-*p*<0.0001; ns, not significant. Primers sequences are listed on Table S21. **(b)** Heatmap depicting expression of *P53* target genes is presented with complete data shown in Table S16 and sorted data shown in Table S19. The color code refers to Z score values. **(c)** Wild-type animals were seeded on EV and *lin-42* RNAi at L1 stage and treated with UV-B (400 and 500 mJ/ cm^2^) after 4h. The % of L1-L2, L3 and L4 stages were counted 44 hours post UV treatment. Significance was measured by an unpaired t-test. n=40-60 in each condition, mean and S.E.M values are presented. *-*p*<0.05; **-*p*<0.01; ***-*p*<0.001; ****-*p*<0.0001; ns, not significant. **(d-f)** Re-analysis of transcriptomic data from cortex of C57BL/6J mice, with samples collected as described in Figure 5a-b. (**d-f**) Markers of DNA damage and DNA repair were analyzed at the beginning (ZT0) and the end (ZT12) of normal sleep (left panel) and during early sleep (NSD) compared to corresponding sleep deprivation points (SD, right panel). Graphs show the expression dynamics of *Cdkn1a* **(d)**, *Gadd45b* **(e)**, and *Ddb2* **(f)**. Significance was measured by unpaired t-test. n= 3-5 in each condition, mean and S.E.M values are presented. *-*p*<0.05; **-*p*<0.01; ***-*p*<0.001; ****-*p*<0.0001; ns, not significant. The complete dataset and the normalization calculation can be found on Table S11-13. **(g)** The model depicting the role of DREAM in mediating the restorative effects of sleep is presented.

## Discussion

In this study, we combined omics, genetics, functional and molecular analyses in three model systems to elucidate the molecular basis of the body-wide functional decline that is caused by a persistent deregulation of the circadian clock and sleep^3,15,16^. As a result, we discovered the conserved chromatin remodeling complex DRM/DREAM to be the central mediator of the sleep and circadian effects at the cellular level. We show that DREAM expression is low during sleep and high during wakefulness, while it fails to be timely lowered in the context of sleep deprivation and clock impairment. High DREAM abundance leads to elevation of histone levels and likely enhanced chromatin compaction. In the context of existing knowledge about DREAM complex, circadian cycle and the protective role of histones and chromatin ^66,67^, our findings suggest that under normal physiological conditions the DREAM-inflicted chromatin change protects the genomic DNA from excessive damage associated with wakefulness ^68,79^.

Conversely, during the sleep phase of the circadian cycle, the chromatin compaction is reversed ^76^ facilitating expression of the genes implicated in the detoxification, homeostasis and repair ranging from translation and splicing to OXPHOS, oxidative stress response and ECM remodeling (Figure 6G). Indeed, many repair activities including autophagy and DNA repair are known to be most effective during sleep ^77,78^, in line with our hypothesis, data and model. In this context, sleep and clock impairments interfere with chromatin re-opening in a DREAM-dependent manner thereby altering repair activities and sensitizing cells to stress and damage to trigger a decline of organ functions. This phenomenon is particularly evident in the organisms predisposed to stress such as nematodes expressing aggregation prone human Aβ or Q40-linked YFP proteins. Because the affected molecular mechanisms are so fundamental, their alteration is likely to affect most cells in an organism, consistent with published observations that circadian/sleep disruptions impair homeostasis of multiple organs^12,44,80^. Our hypothesis is also consistent with the large body of reports linking sleep deprivation to human diseases and inferior organismal responses to stressors ranging from infections and cognitive challenges to exercise and diet^81–84^. We therefore lay out for the first time a mechanistic model that integrates and explains most of the current knowledge about organismal and cellular effects of disrupted sleep and circadian dysfunction. From the translational standpoint, we additionally show that pharmacological and genetic inhibition of DREAM can possibly mimic the restorative effects of sleep at the cellular level.

Notably, our findings suggest that the DREAM complex simultaneously regulates multiple cellular functions via controlling the abundance of histones and (likely) the accessibility of the respective chromatin regions. This model is supported by the previous report detecting high abundance of the histone variant HTZ-1/H2A.Z in the gene body of DRM/DREAM-repressed cell cycle targets^29^, with HTZ-1/H2A.Z being one of the principle histones we found to cooperate with DRM/DREAM in de-regulating the cellular resilience of clock-distorted animals. Consistently, only a minor fraction of the molecular targets altered by clock disruption in a DREAM-dependent manner harbors relevant DREAM binding sites in their promoter regions (Table S10), supporting an indirect regulation by modulation of the chromatin structure. Interestingly, this histone-dependent mode of DREAM action is distinct from the well-documented capacity of DRM/DREAM to repress multiple pathways implicated in DNA repair^72,73,85,86^, which relies on the direct binding of DREAM to the regulatory elements of affected genes. Notably, also the predominantly implicated DRM/DREAM subunits differ between DNA repair and circadian functions of DRM/DREAM with *lin-52*, *dpl-1*, *efl-1* and *lin-35* having the strongest impact in DNA repair^72^ and *lin-9*, *lin-54* and *lin-53* regulating the consequences of the circadian disruption. Our findings thus support the likely existence of two different modes of how specific components of the DREAM complex regulate complex gene expression networks: (a) via direct binding and recruitment of co-factors and/or co-repressors^31,32^ and (b) via changes in histone composition and chromatin structure (^29^ and this study). The dependency of the circadian response on the presence of particular DREAM subunits opens a possibility of targeted interventions that modulate the “sleep-specific” role of DREAM without interfering with its activities in other cellular processes. An additional path of mitigating potential side effects derives from our new mechanistic model presented in Figure 6G: we propose that the administration of DREAM inhibitors shall be restricted to the time designated to sleep as opposed to the 24h-long chronic administration, thus replicating the natural circadian fluctuations of DREAM activity. In follow-up studies, we anticipate to functionally test these hypotheses *in vivo* in vertebrates.

## Supporting information

Statistics source data

## Acknowledgements

We thank the Proteomics Core Facility at FLI, and especially Joanna Kirkpatrick, Erika Kelmer Sacramento, Emilio Cirri, and Therese Dau for supporting this study. The Core Facility Next Generation Sequencing (Ivonne Goerlich, Marco Groth) of the FLI is gratefully acknowledged for their technological support in library preparation and mRNA sequencing. Marcus Wildt for his support in data structure and coding. The FLI is a member of the Leibniz Association and is financially supported by the Federal Government of Germany and the State of Thuringia. ISV was supported by the doctoral fellowship from the Leibniz Graduate School on Aging (LGSA). ME was supported by the EU-ESF Thüringer Aufbaubank funding (2019 FGR 0082). ME is currently funded by the Carl-Zeiss-Stiftung via IMPULS consortium and is a member of Cluster of Excellence Balance of the Microverse funded by the Deutsche Forschungsgemeinschaft (DFG). ME is also funded by the ERC CoG LifeLongFit of the European Commission. *C. elegans* strains were obtained from the Caenorhabditis Genetics Center (CGC), which is funded by NIH. FJRE is supported by a contract of the VPPI-University of Sevilla (Contratos de Personal Investigador en Formación). Work in the Olmedo lab is supported by the project PID2022-139009OB-I00, funded by MCIN/AEI/10.13039/501100011033 and FEDER, UE.

## Author contributions

ME and ISV conceptualized and designed the study; ISV, FJRE, KS, MB, AT and LL performed experiments; ME, MO, MF, SH and OE supervised data analysis and visualization; ME, ISV, FJRE, KS, MB, MF, KR, SH, MO and OE analyzed the data; ISV prepared figures and tables, and performed statistical analyses; ME wrote the manuscript, ISV, MF, SH, MO and OE reviewed the manuscript.

## Data availability

The mass spectrometry proteomics data have been deposited to the ProteomeXchange Consortium via the PRIDE (Perez-Riverol et al., 2019) and MassIVE repositories with the respective dataset identifiers PXD024273 and MSV000091972. The RNA-seq data have been deposited to the NCBI Gene Expression Omnibus (GEO) with data set identifier GSE250018.

## Animal experiments

Experiments in mice comply with all relevant ethical and experimental regulations, as validated by the dedicated authority of the state of Thuringia (Germany).

---

**Fig. S1.**
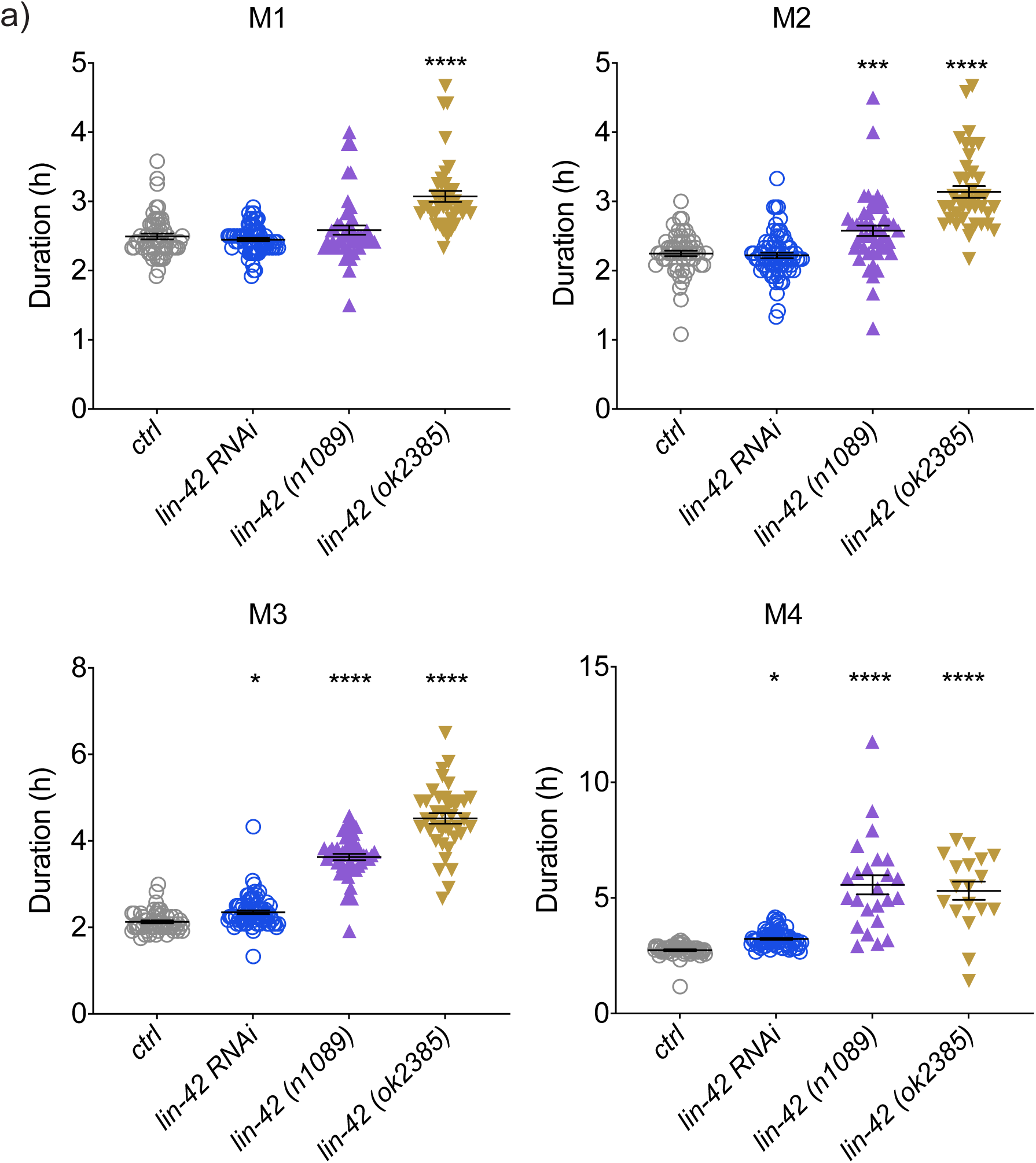
RNAi-mediated inactivation of *lin-42*/PER interferes with clock-dependent developmental timing. **(a)** Developmental timing (duration of molting stages M1-M4) of control (ctrl), *lin-42* RNAi knock-down, *lin-42(n1089)* mutant and *lin-42(ok2385)* mutant animals is shown. Control (EV RNAi) and *lin-42* RNAi cohorts were established in the wild type (N2 Bristol strain) background, *lin-42(n1089)* mutant and *lin-42(ok2385)* mutant animals were fed with EV RNAi, and all worms were fed with D-Luciferin. Molting stage durations were quantified at 20°C based on relevant luminescence signals. Each dot represents one animal; average and S.E.M values are plotted. n=5-20 worms per condition in each trial, and results of 4 independent trials are combined. One-way ANOVA was used for the statistical assessment, two-tailed *p* values were computed *-*p*<0.05; **-*p*<0.01; ***-*p*<0.001; ****-*p*<0.0001.

**Fig. S2.**
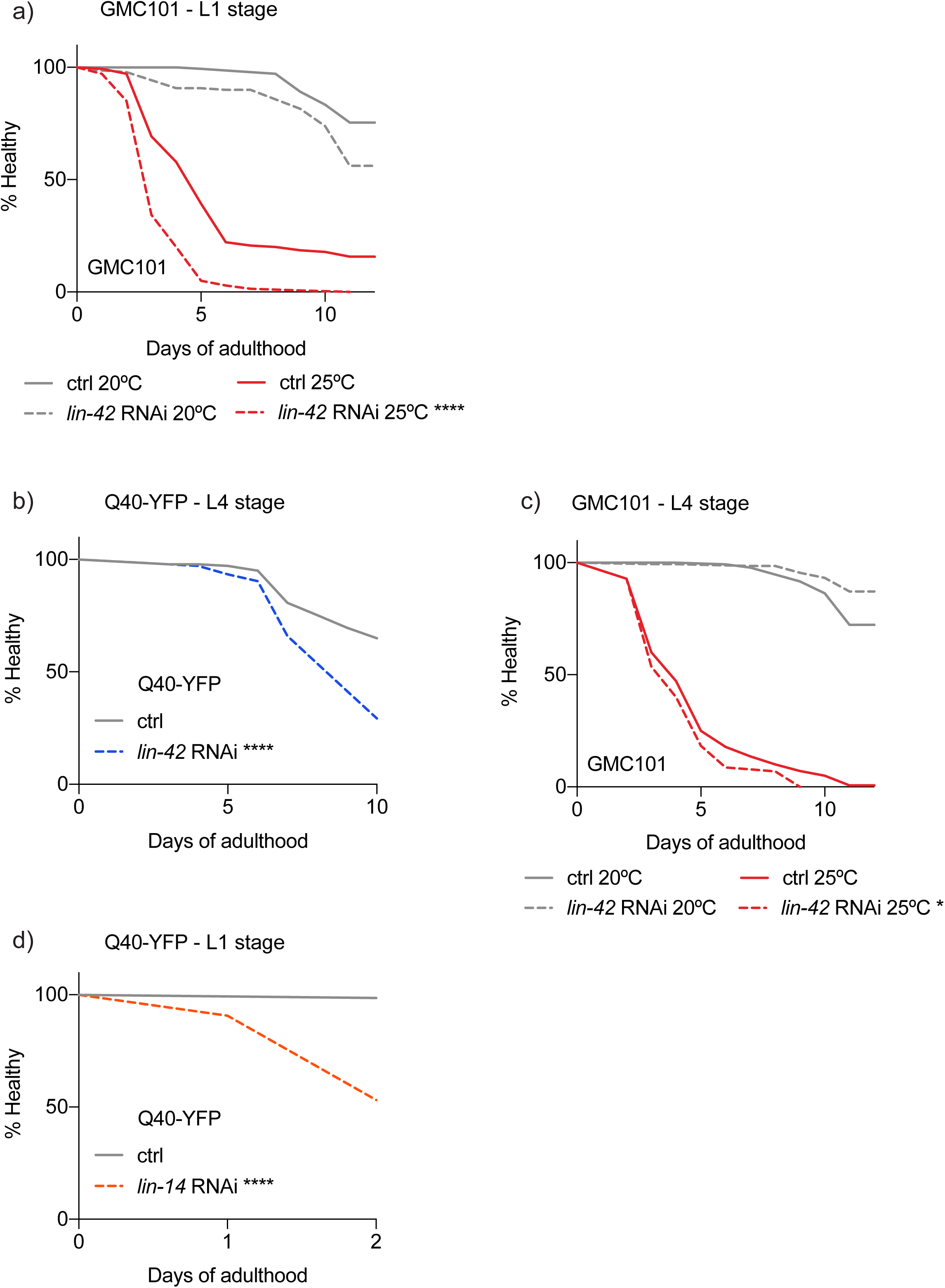
Clock disruption exacerbates muscle paralysis in *C. elegans* models of Alzheimer’s and Huntington’s diseases. **(a)** *unc-54*p::human Aβ_1-42_ worms (GMC101 strain) were age-synchronized and treated with EV (ctrl) or *lin-42* RNAi from the L1 stage. Until the L4 stage the worms were cultivated at 20°C and then shifted to 25°C, to trigger amyloid-beta toxicity. **(b)** Age-synchronized population of *unc-54*p::Q40::YFP animals were fed with EV (ctrl) or *lin-42* RNAi from the L4 stage. **(c)** *unc-54*p::human Aβ_1-42_ worms were age-synchronized and treated with EV (ctrl) or *lin-42* RNAi from the L4 stage. The shift to 25°C was performed as described in (a). **(d)** Age-synchronized population of *unc-54*p::Q40::YFP animals were fed with EV (ctrl) or *lin-14* RNAi from the L1 stage. Paralysis was scored daily in all cases (**a-d**). n=140 worms per condition, graphs show representative results of at least three independent experiments. Significance was measured by the Log-rank Mantel-Cox test, two-tailed *p* values were computed. *-*p*<0.05; ****-*p*<0.0001, ns, not significant.

**Fig. S3.**
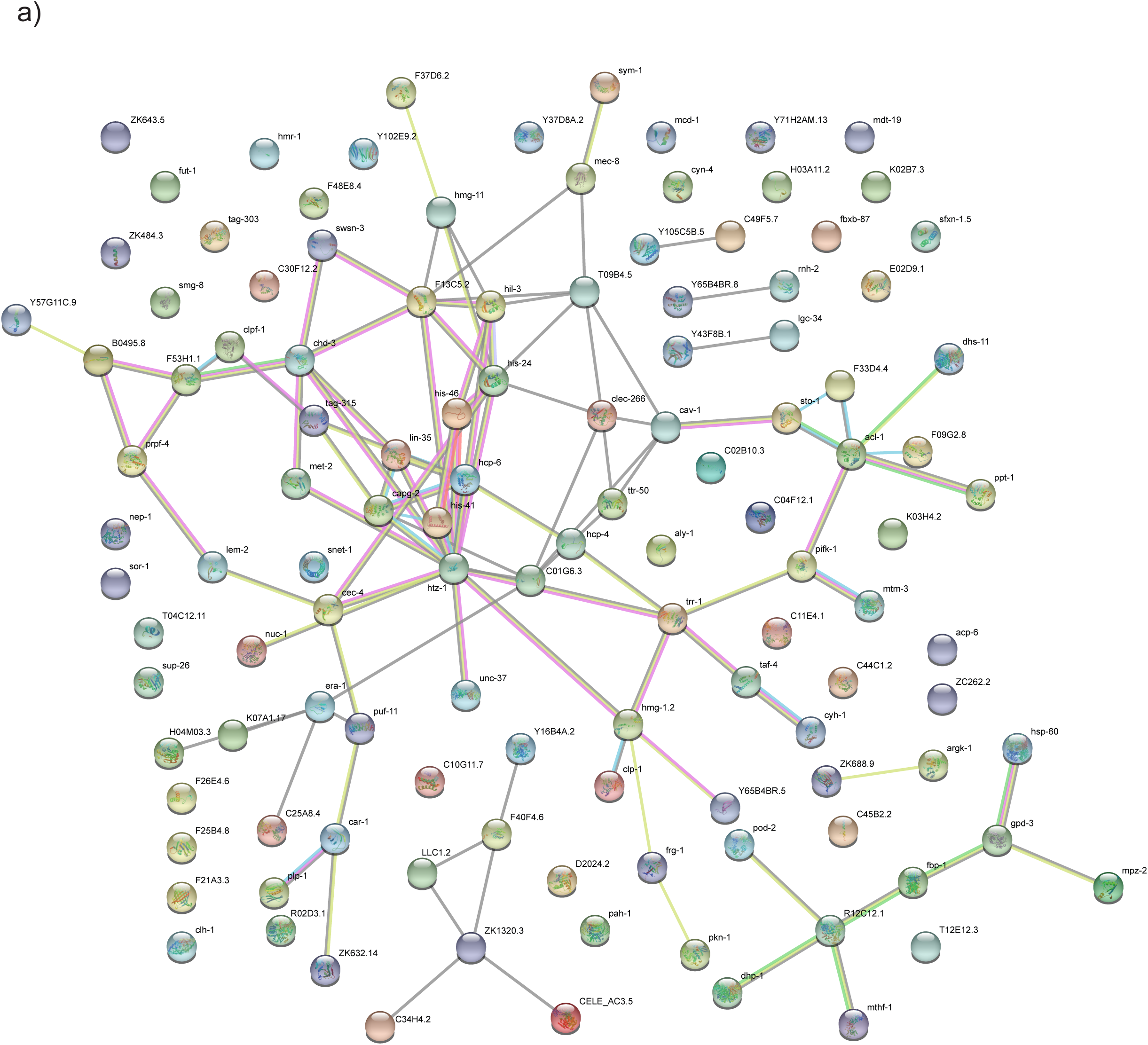
Protein-protein interaction analysis reveals DRM/DREAM complex as a putative central target of clock impairment. STRING analysis was performed using version 11 (latest version at the time of the analysis) with the protein groups that contributed strongest to the PC1 separation in Fig. 2b (the complete list and data can be found in Table S4). The absolute rotational weight of the included proteins had a cutoff of >0.023, and 120 proteins had a significant network enrichment of *p*<1.0e^−16^ (Table S5). Colored network nodes represent query proteins. Edges represent meaningful protein-protein associations. Turquoise and purple edges represent known interactions, with purple depicting experimentally tested interactions. Green, red and blue edges represent predicted interactions, with green representing gene neighborhood, red – gene fusions, and blue – gene co-occurrence. Light green edges represent results of text mining, black edges – co-expression, and light blue edges – protein homology.

**Fig. S4.**
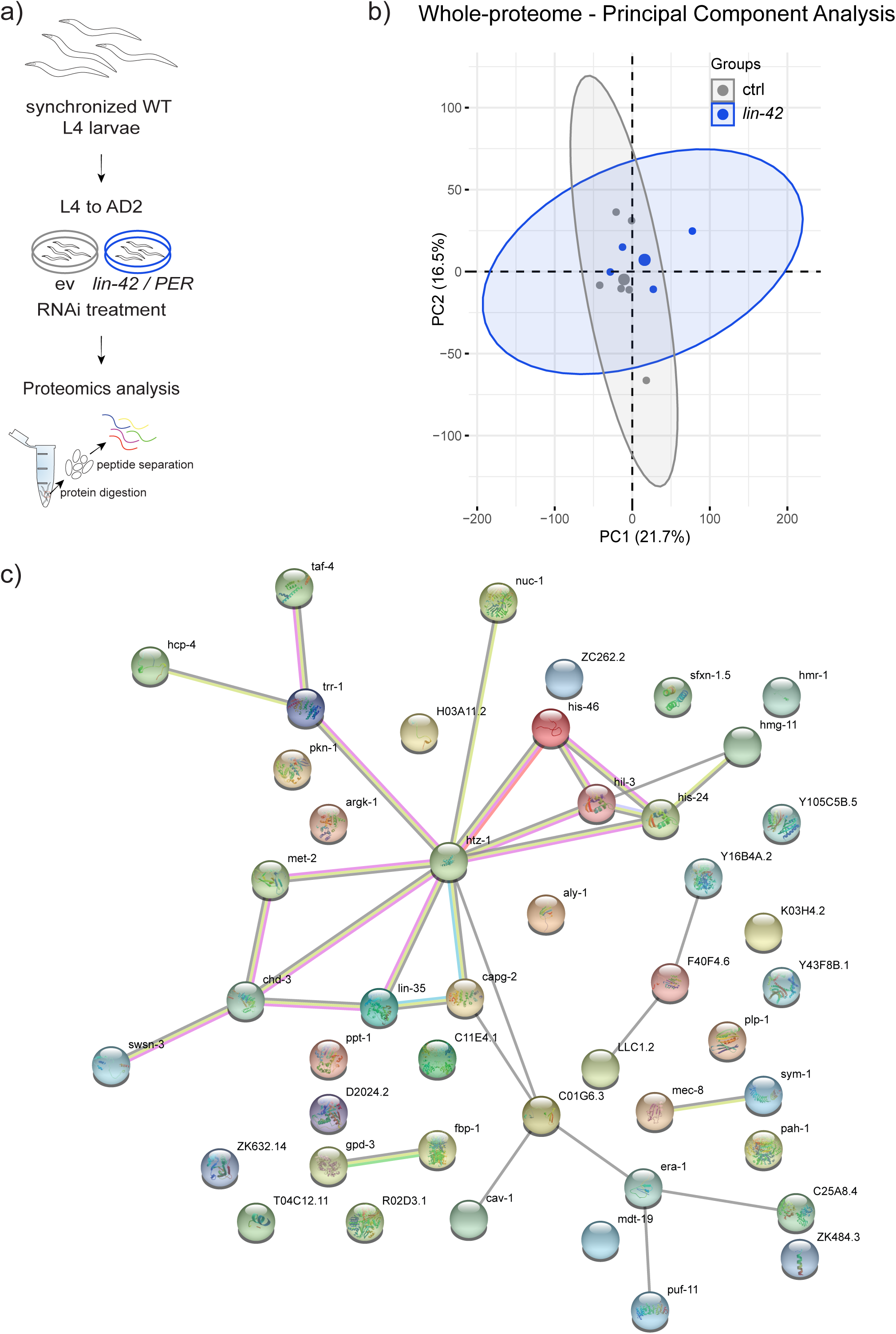
Whole proteome changes are comparable between animals subjected to *lin-42*/PER inactivation during development and adulthood. **(a)** Experimental design. **(b)** Whole-proteome principal component analysis of wild-type *C. elegans* treated with EV (ctrl) or *lin-42* RNAi from the L4 stage is shown. Small dots are representative of individual replicas, n=800 worms per sample. **(c)** STRING analysis using version 11 (latest version at the time of the analysis) was performed with the protein groups that contributed the strongest to the PC1 separation in Fig. S4b. The related source data can be found in Tables S2 (complete data) and S4 (STRING data). The absolute rotational weight of the proteins had a cutoff of >0.01, and 46 proteins had a significant network enrichment of *p*<9.18e^−12^ (Table S5). The color code of nodes and edges is as explained in Figure S3.

**Fig. S5.**
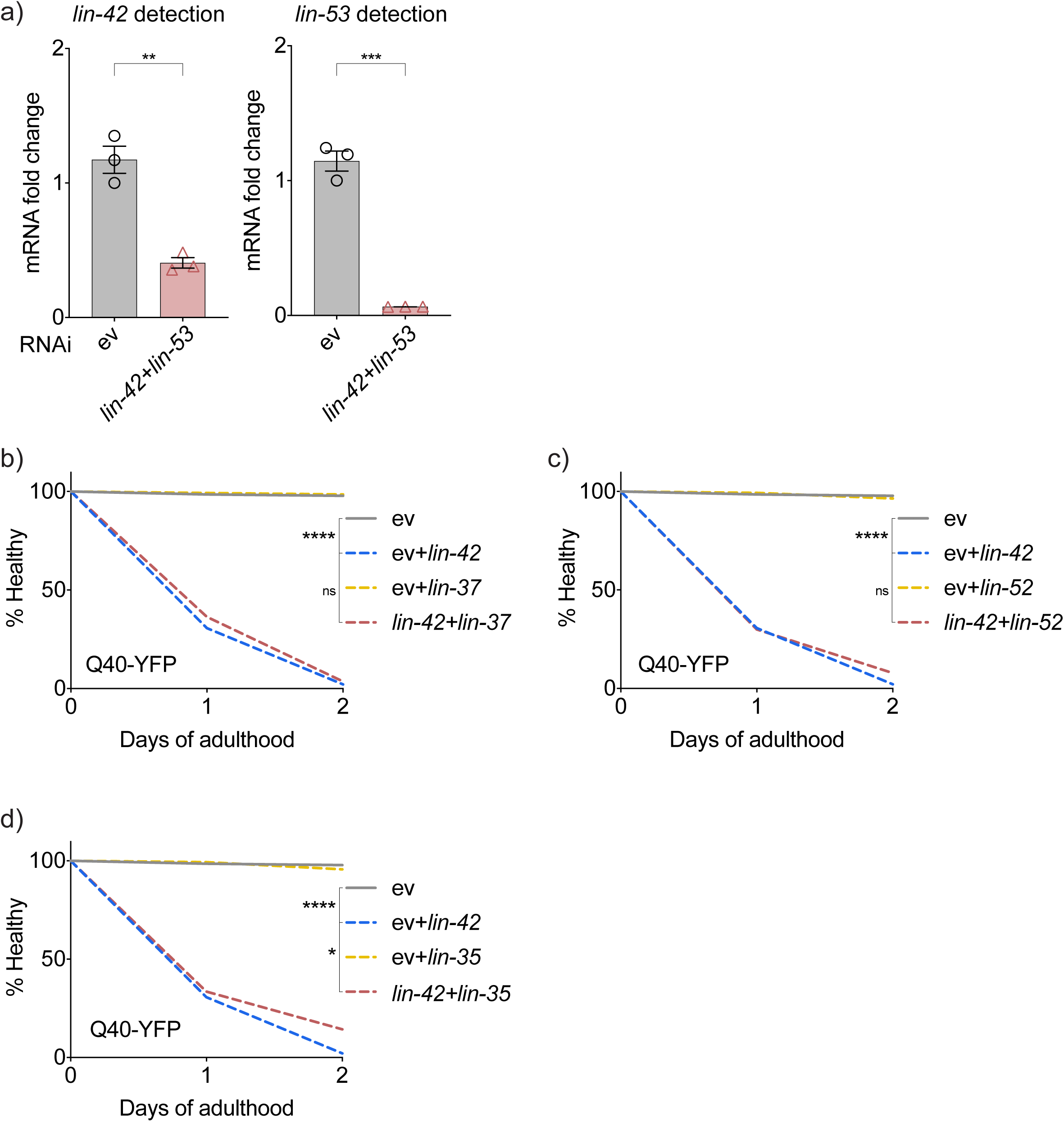
The interplay between DRM/DREAM and clock impairment demonstrates subunit specificity. **(a)** 250 worms per condition were age-synchronized and treated with indicated RNAi combinations from the L1 stage, and *lin-42* and *lin-53* gene expression was measured by RT-qPCR on AD2. Graphs are representative of three independent experiments. Mean and S.E.M. values are presented. mRNA expression fold change was calculated by the ΔΔCt method. Significance was measured by unpaired t-test, two-tailed *p* values were computed. **-*p*<0.01; ***-*p*<0.001; ns, not significant. Primers sequences are listed on Table S20. **(b-d)** *unc-54*p::Q40::YFP animals were age synchronized and fed from the L1 stage with EV or *lin-42* RNAi and in combination with RNAi against specific DREAM subunits or interactors: *lin-37* **(b)**, *lin-52* **(c)**, *lin-35* **(d)**. Paralysis was scored daily. n=140 worms per condition, graphs are representative of at least three independent experiments. Significance was measured by the Log-rank Mantel-Cox test, two-tailed *p* values were computed. ****-*p*<0.0001; *-*p*<0.05, ns, not significant.

**Fig. S6.**
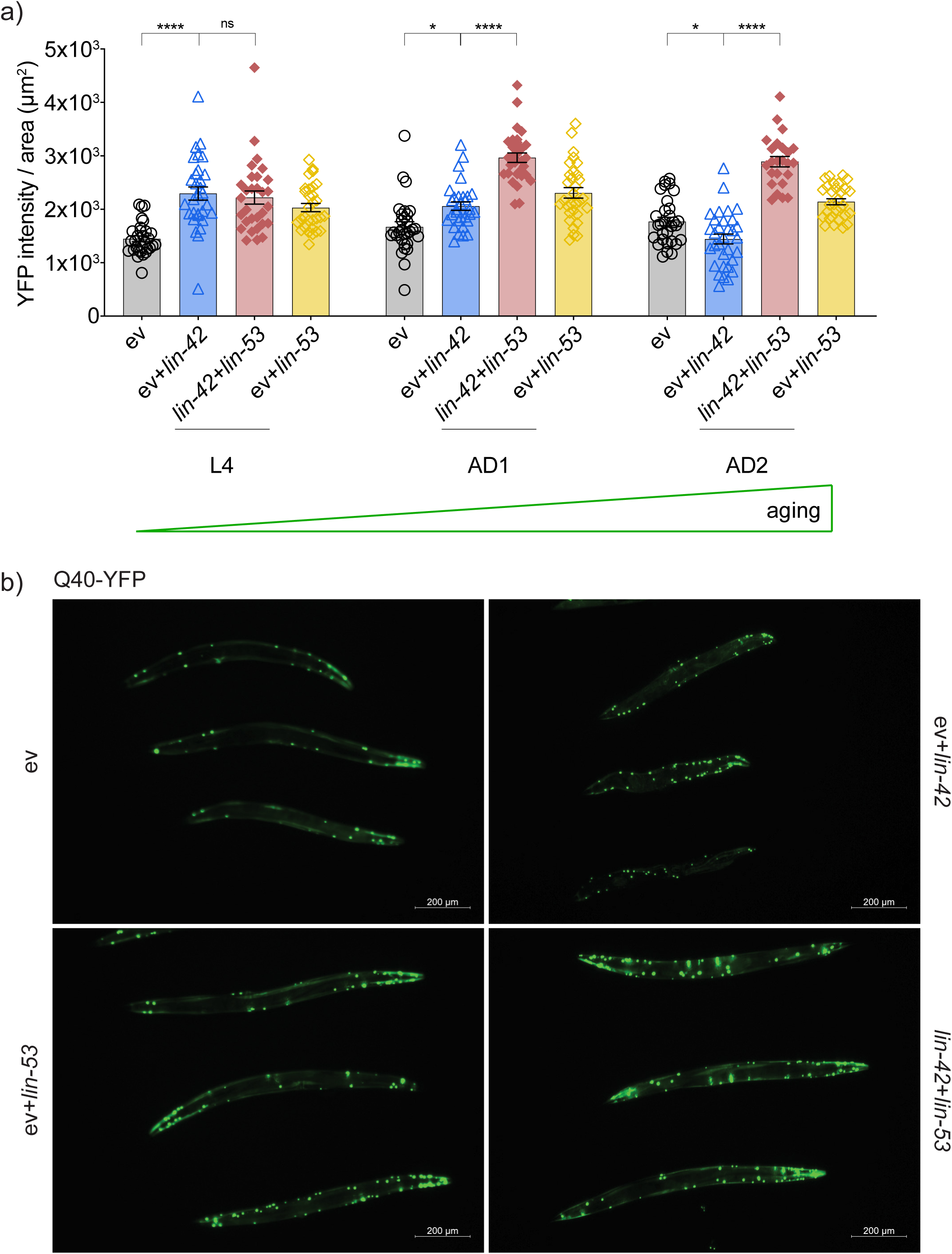
*lin-53*/DREAM inactivation restores protein homeostasis in clock-disrupted animals expressing polyglutamine-linked YFP. *unc-54*p::Q40::YFP worms were age-synchronized and fed with indicated RNAi combinations from the L1 stage and imaged at three different time points: L4, AD1, and AD2; EV – empty vector control. **(a)** YFP intensity per total body area is shown. n=26-32 worms per condition, the graph is representative of three individual experiments. Error bars are S.E.M. Significance was measured by one-way ANOVA within each age group separately and using Sidak’s multiple comparison test, two-tailed *p* values were computed. *-*p*<0.05; ****-*p*<0.0001; ns, not significant. **(b)** Representative images of AD2 *unc-54p*::Q40::YFP animals treated with indicated RNAi combinations. Worms were imaged in an AxioZoom v.16 microscope with 90x magnification and exposure time of 150 ms for YFP and 4.8 ms for brightfield imaging. The scale bar is 200 µm.

**Fig. S7.**
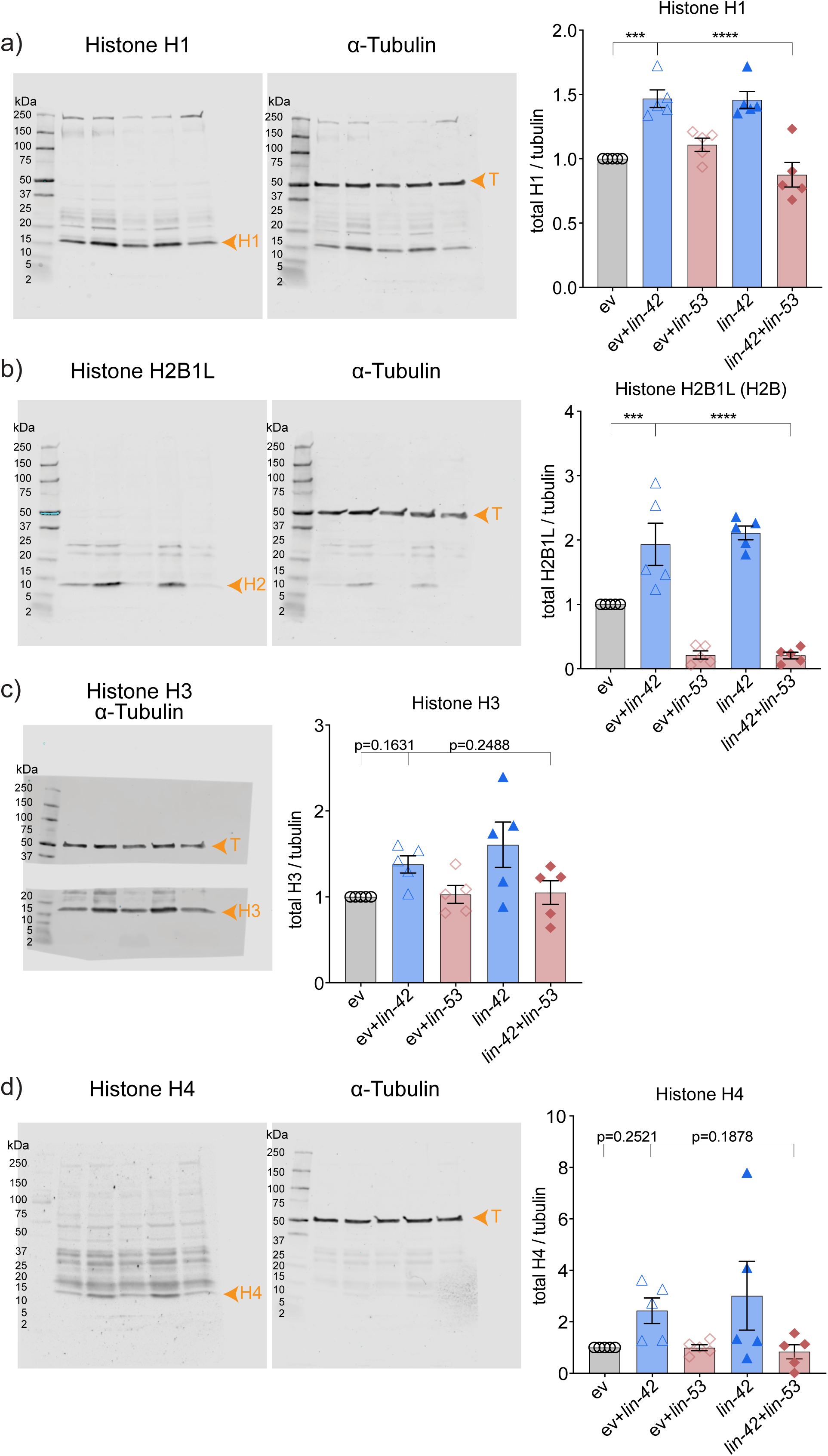
Clock disruption enhances the expression of histones in a DRM/DREAM-dependent manner. **(a-d, left panels)** Full gel scans depict expression of indicated histones using tubulin as loading control. **(a)** H1, **(b)** H2B1L, **(c)** H3, **(d)** H4. Size-consistent bands used for quantification are marked by arrowheads. H= histone and T= tubulin. **(a-d, right panels)** The corresponding quantification graphs are presented. Quantification of 5 independent experiments represented by individual data points is shown; n=800 worms in each condition. Error bars are S.E.M. Significance was measured by one-way ANOVA, two-tailed *p* values were computed. ***-*p*<0.001, ****-*p*<0.0001, ns, not significant.

**Fig. S8.**
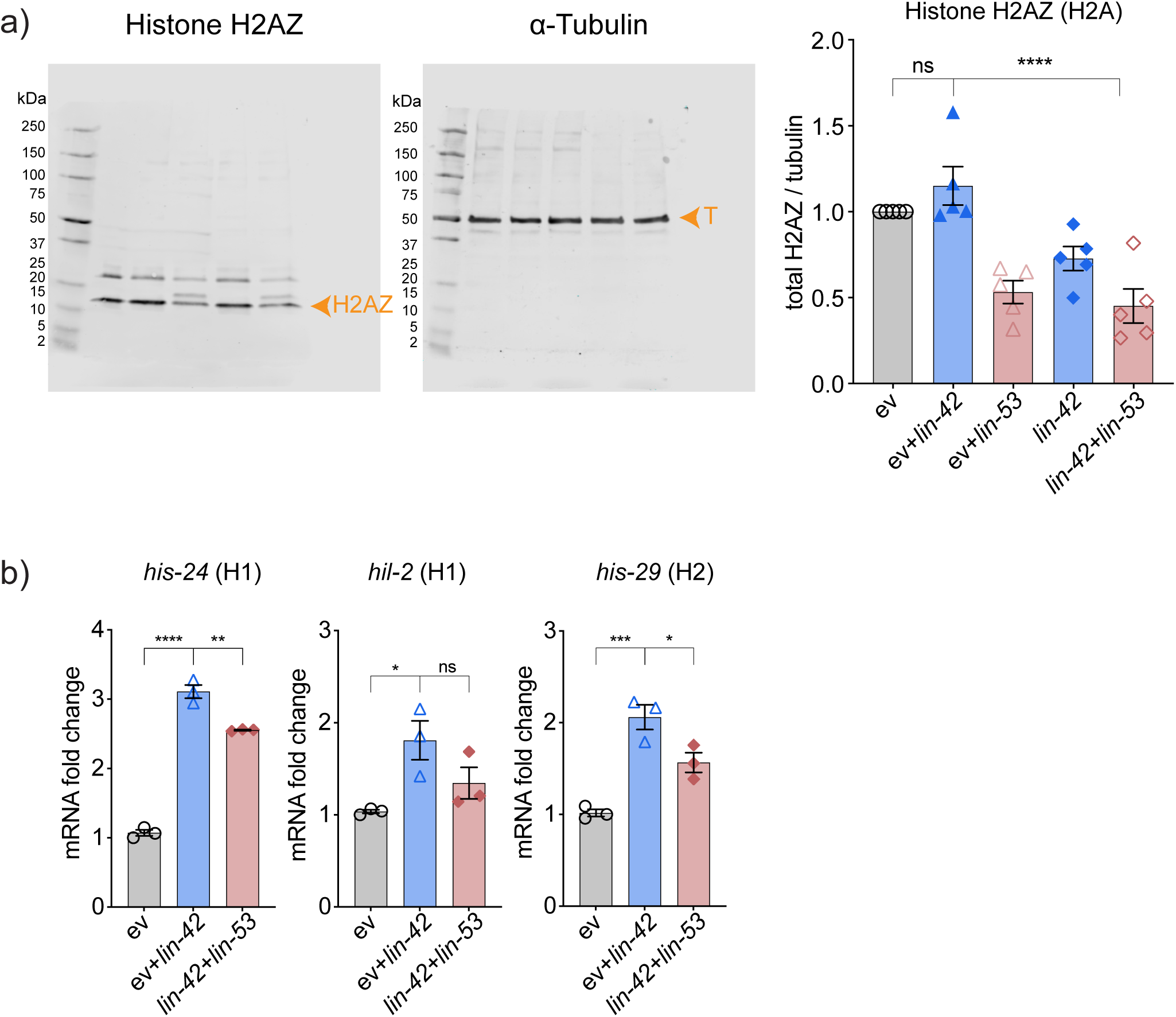
Clock disruption elevates the expression of histones in a DRM/DREAM-dependent manner. **(a)** Worms were treated and analyzed as described in Figure 4a-b, using antibody specific to histone H2AZ. Representative full gel scans (left) and respective quantification (right) are shown. The graph summarizes the results of five independent experiments. n=800 per condition and error bars are S.E.M. Significance was measured by one-way ANOVA, two-tailed *p* values were computed. ****-*p*<0.0001, ns, not significant. **(b)** Worms were treated as in **(a)** and mRNA expression levels of *his-24* (H1), *hil-2* (H1), and *his-29* (H2) were assessed by RT-qPCR. n=250 worms per condition, graphs are representative of three independent experiments. Error bars are S.E.M.; mRNA fold change was calculated by the ΔΔ Ct method. Significance was measured by one-way ANOVA within each group separately and applying Sidak’s multiple comparisons test, two-tailed p values were computed. *-p<0.05; **-p<0.01; ***-p<0.001; ****-p<0.0001; ns, not significant. Primers sequences are listed on Table S20.

**Fig. S9.**
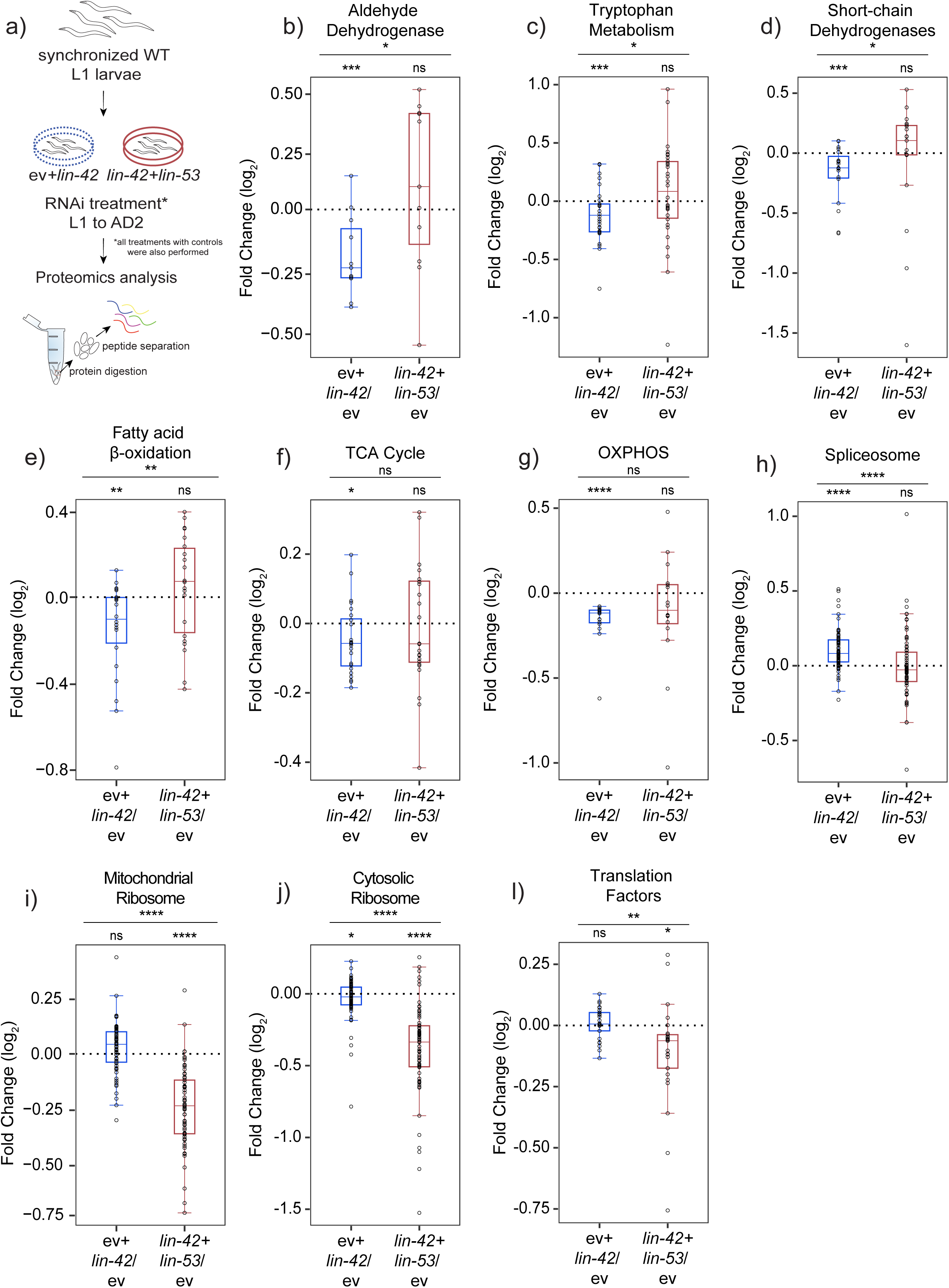
Circadian clock disruption interferes with fundamental cellular processes in a DRM/DREAM-dependent manner. **(a-l)** Animals were treated and analyzed as described in Figure 4c-h. Box plots show the relative expression of proteins involved in aldehyde dehydrogenase **(b)**, tryptophan metabolism **(c)**, short-chain dehydrogenases **(d)**, fatty acid β-oxidation **(e)**, TCA cycle **(f)**, OXPHOS **(g)**, spliceosome **(h)**, mitochondrial ribosome **(i)**, cytosolic ribosome **(j)**, and translation factors **(l)**. Individual proteins are shown as dots, the median fold change of each group is shown as a horizontal line; the upper and lower limits of the boxplot indicate the first and third quartile and the whiskers extend 1.5 times the interquartile range from the limits of each box. n=800 per condition and 4 independent populations were measured. Within each box plot, the significance was assessed by the Mann-Whitney Wilcoxon rank-sum test, and Wilcoxon test was used for the comparison between the two sets of fold changes. Two-tailed *p* values were computed in all cases. *-*p*<0.05; **-*p*<0.01; ***-*p*<0.001; ****-*p*<0.0001. The lists and data of all proteins can be found in Tables S8 (complete data) and S9 (box plot data).

**Fig. S10.**
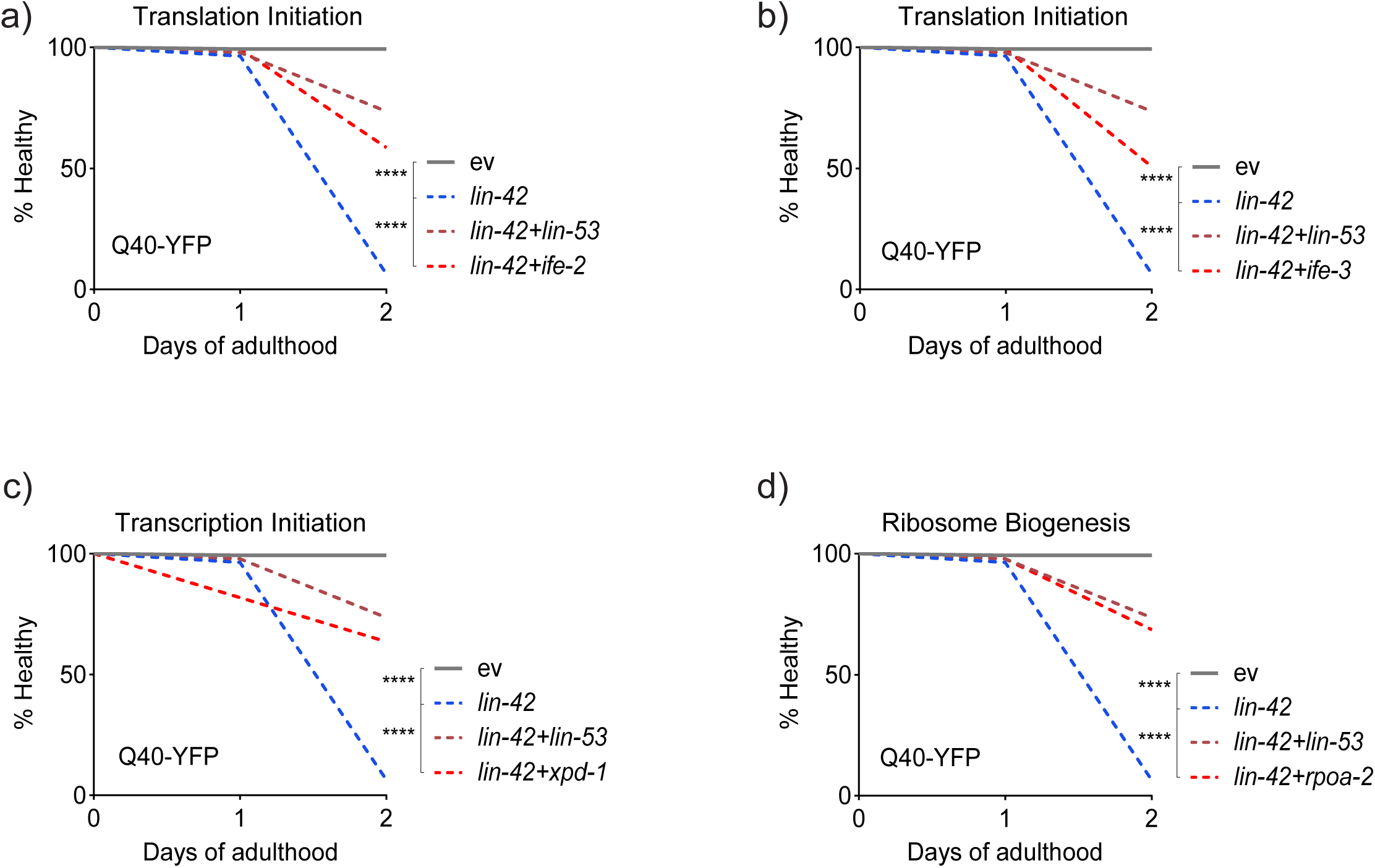
DREAM-dependent pathways are functional drivers of the negative effects of circadian disruption. **(a-d)** Age-synchronized *unc-54*p::Q40::YFP animals were fed with EV or *lin-42* RNAi in combination with RNAis targeting specific genes implicated in the clock/DREAM co-dependent pathways: translation initiation, ribosome biogenesis and transcription initiation, from the L1 stage, paralysis was scored daily. Co-targeting with *ife-2* **(a)**, *ife-3* **(b)**, *xpd-1* **(c),** and *rpoa-2* **(d)** is shown. n=140 worms per condition, graphs are representative of at least three independent experiments. Significance was measured by the Log-rank Mantel-Cox test, two-tailed *p* values were computed. ****-*p*<0.0001.

**Fig. S11.**
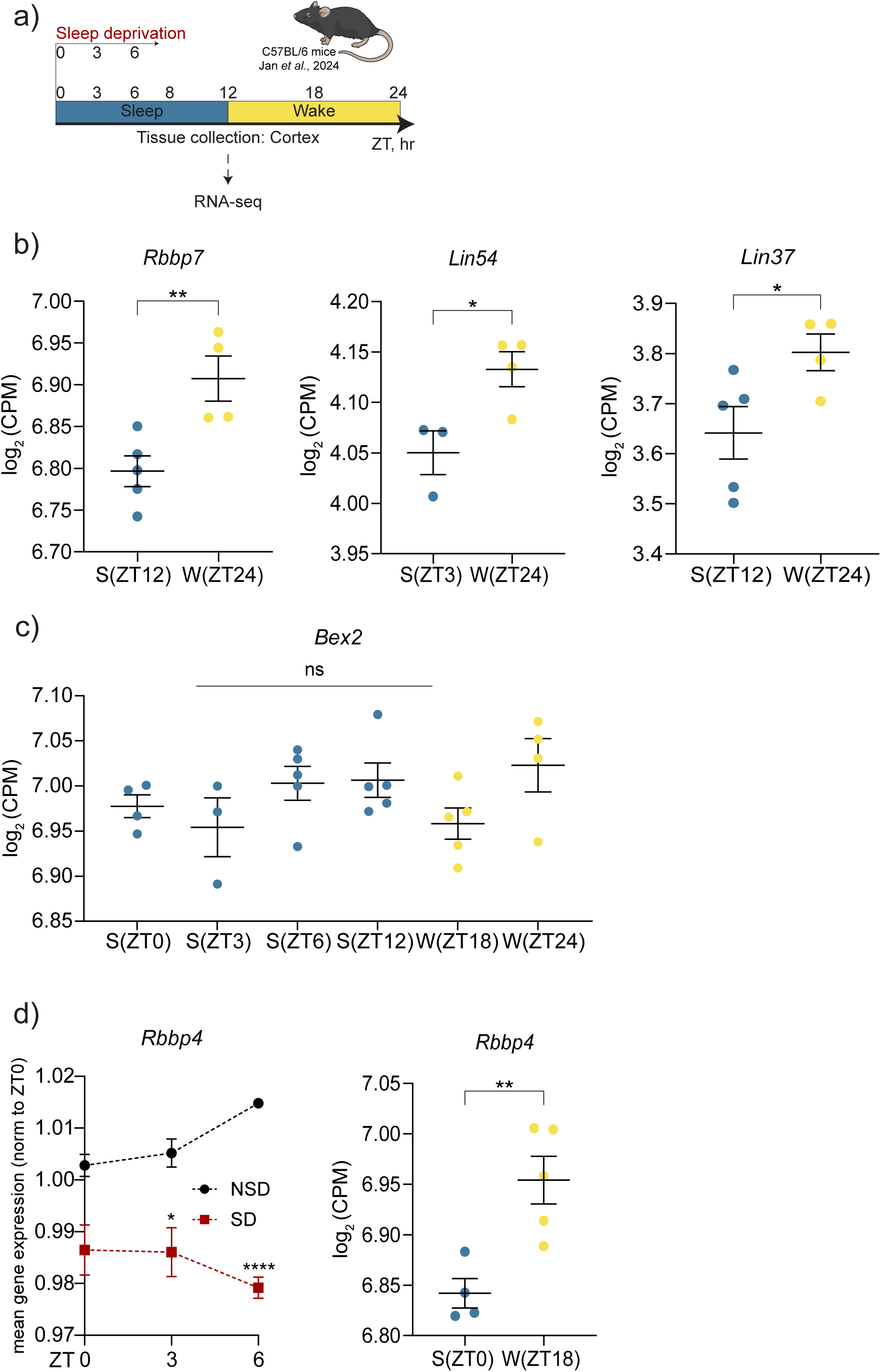
DREAM expression changes during the natural sleep-wake cycle. **(a)** Experimental design of Jan *et al.*, 2024 is depicted. **(b-d)** The transcriptomic data was analyzed as in Fig. 5a-c. **(b)** Expression of DREAM subunits *Rbbp7*, *Lin54*, and *Lin37* during normal sleep (S, blue circles) and wakefulness (W, yellow circles) is shown as Log_2_ CPM, each dot represents and independent sample. n= 3-5 in each condition, mean and S.E.M values are presented. Significance was measured by unpaired t-test at indicated ZTs. *-*p*<0.05; **-*p*<0.01; ns, not significant. **(c)** The expression of a non-cycling gene *Bex2* is depicted at all time points of sample collection as a control. **(d)** The expression of the *Rbbp4* DREAM subunit during sleep and wakefulness (right panel) and comparing early sleep (NSD) to sleep deprivation (SD, left panel) is shown. In **(d, left panel**) significance was measured by unpaired t-test. n= 3-5 in each condition, mean and S.E.M values are presented. *-*p*<0.05; **-*p*<0.01; ***-*p*<0.001; ****-*p*<0.0001; ns, not significant. The complete dataset and the normalization calculation can be found on Table S11 and S12.

**Fig. S12.**
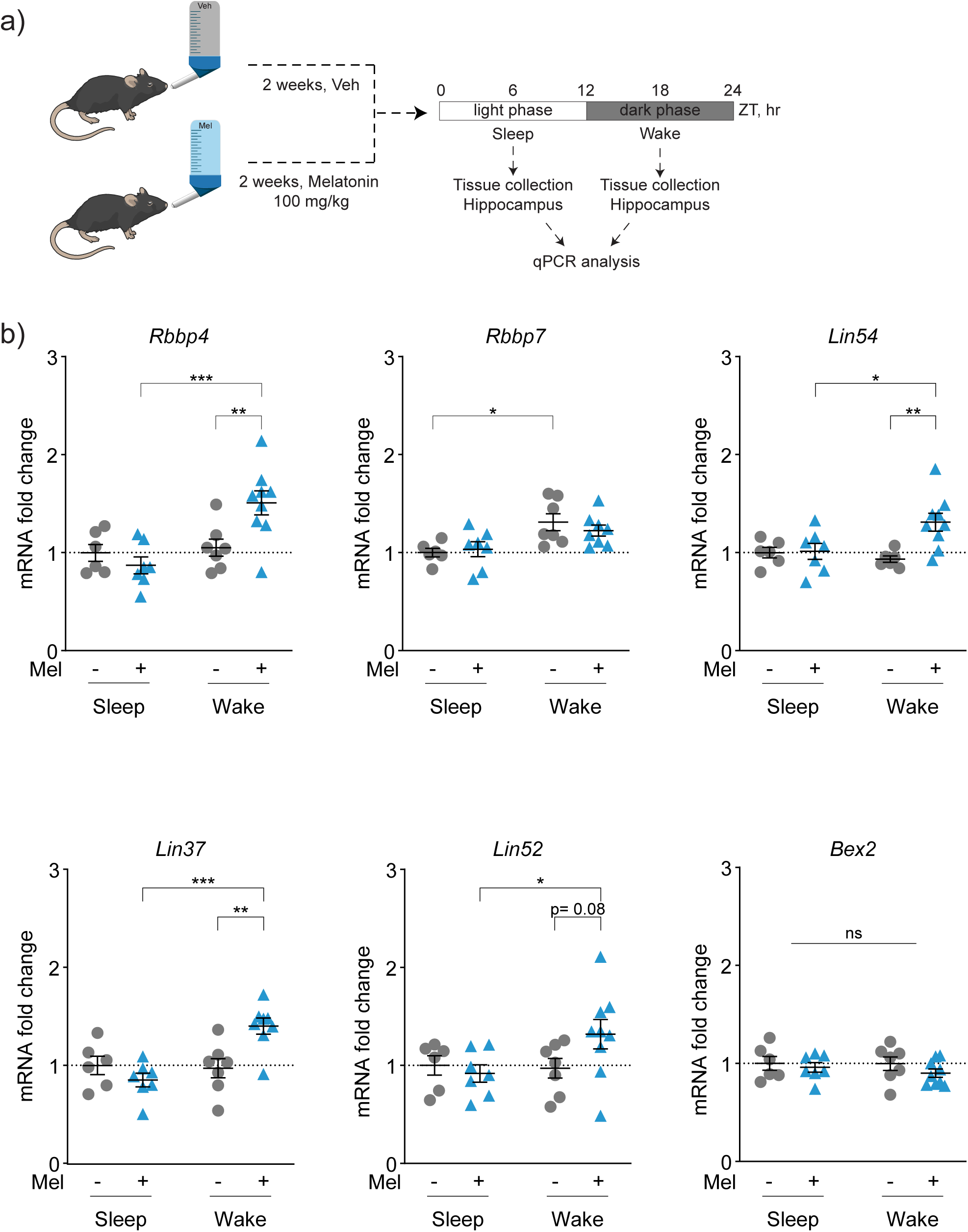
The amplitude of circadian DREAM oscillations is enhanced by Melatonin treatment. **(a)** Experimental design. **(b)** Wild-type mice were treated with melatonin (100 mg/kg) or vehicle for 2 weeks. Hippocampi were collected at both light (sleep) and dark (sleep) phases and samples processed for RT-qPCR analysis. The expression analysis of DREAM subunits *Rbbp4*, *Rbbp7*, *Lin54*, *Lin37*, *Lin52* and the non-cycling gene *Bex2* were performed by RT-qPCR. Data is shown as mRNA expression fold changes calculated by the ΔΔCt method. Significance was measured by multiple comparisons two-way ANOVA with a Bonferroni correction. n= 6-9 in each condition, mean and S.E.M values are presented. *-*p*<0.05; **-*p*<0.01; ***-*p*<0.001; ****-*p*<0.0001; ns, not significant. Primers sequences are listed on Table S22.

**Fig. S13.**
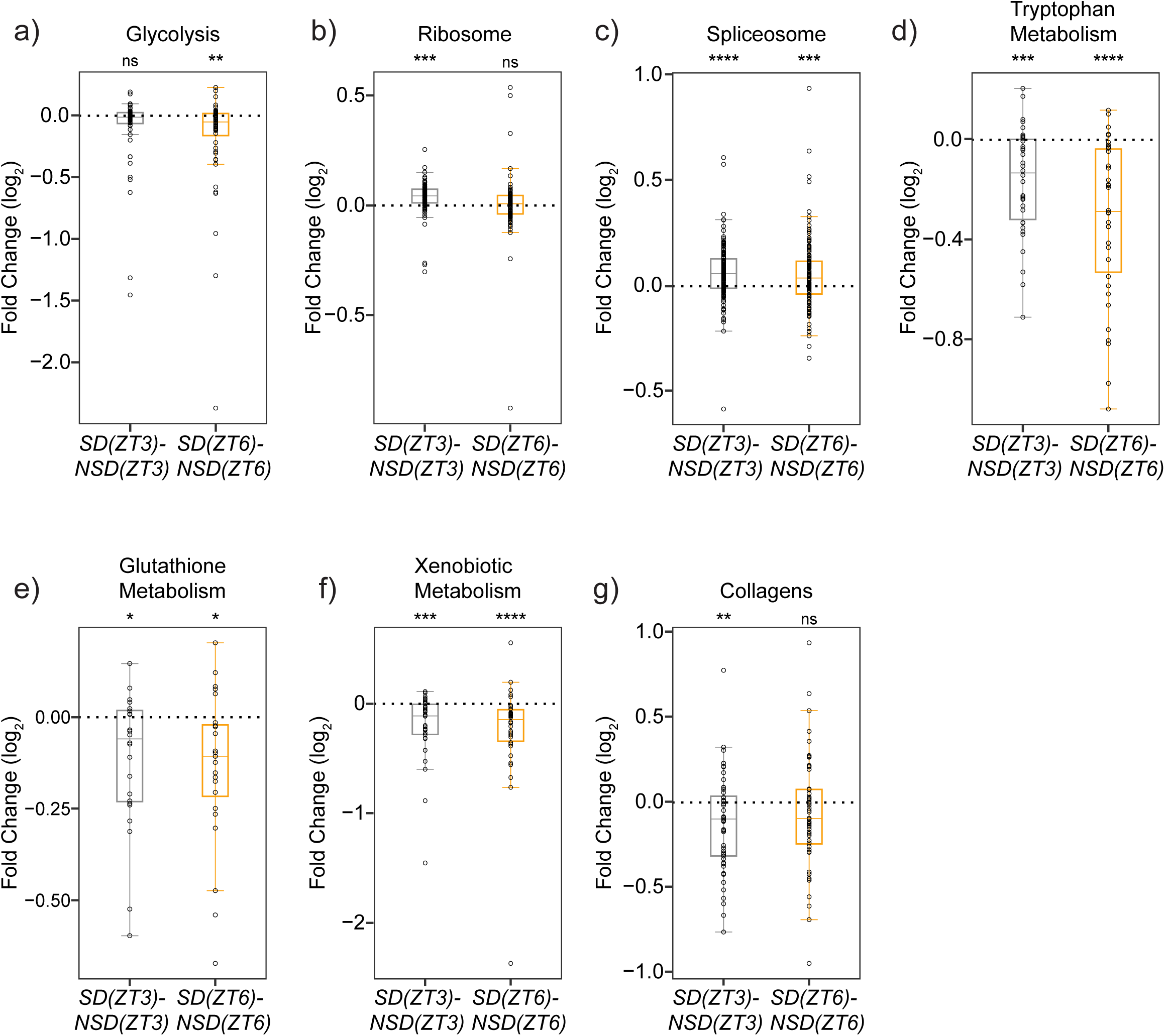
Same fundamental cellular processes are altered by sleep disruption in mice as in nematodes. **(a-g)** Transcriptomics data analyzed as in Fig. 5d. Box plots showing relative expression of glycolysis **(a)**, ribosome **(b)**, spliceosome **(c)**, tryptophan metabolism **(d)**, glutathione metabolism **(e)**, xenobiotic metabolism **(f)** and collagens **(g)** genes are presented. Individual genes are shown as dots, the median fold change of each group is shown as a horizontal line; the upper and lower limits of the boxplot indicate the first and third quartile and the whiskers extend 1.5 times the interquartile range from the limits of each box. n=6-7 mice per condition. Within each box plot, the significance was assessed by the Mann-Whitney Wilcoxon rank-sum test, and the Wilcoxon test was used for the comparison between the two sets of fold changes. Two-tailed *p* values were computed in all cases. *-*p*<0.05; **-*p*<0.01; ***-*p*<0.001; ****-*p*<0.0001. The lists and data of all genes can be found in Tables S13 (complete data) and S15 (box plot data).

**Fig. S14.**
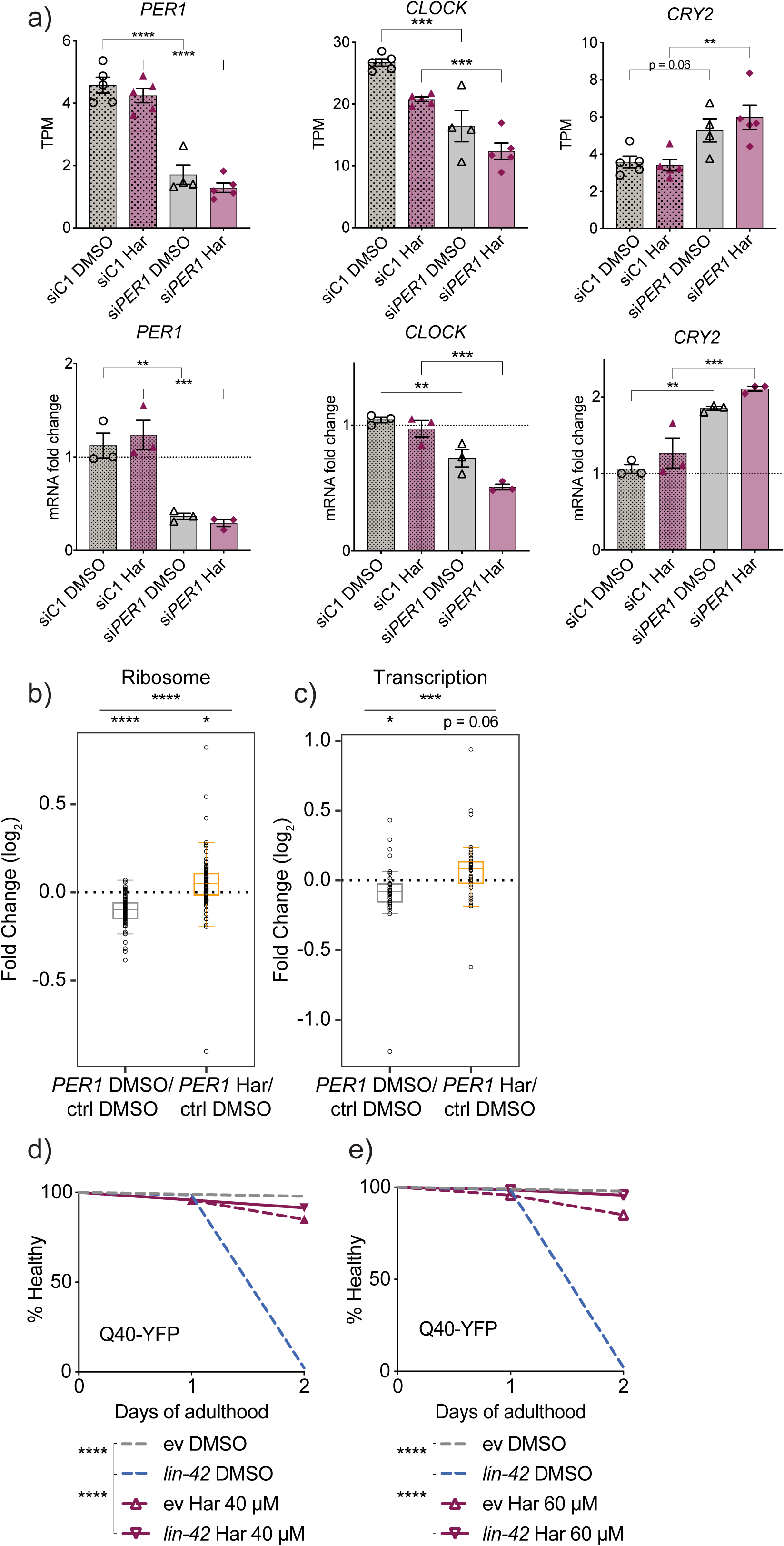
Same fundamental cellular activities are regulated by clock/DREAM axis in human cells as in nematodes. **(a)** Human RPE cells were treated and analyzed as described in Fig. 6a, n=4-5 independent cultures per condition. Expression analysis of *PER1*, *CLOCK* and *CRY2* genes are shown. RT-qPCR (lower panels) and transcriptomic measurements (upper panels) are presented as described in Fig. 6a. Significance was measured by one-way ANOVA within each group separately and applying Sidak’s multiple comparisons test, two-tailed *p* values were computed. ****-*p*<0.0001. Primers sequences are listed on Table S21. **(b-c)** Transcriptomic data was analyzed as in Fig. 5f. Box plots showing relative expression of the ribosome **(b)**, and RNA polymerase (transcription) **(c)** genes are presented. Individual genes are shown as dots, the median fold change of each group is shown as a horizontal line; the upper and lower limits of the boxplot indicate the first and third quartile and the whiskers extend 1.5 times the interquartile range from the limits of each box. Within each box plot, the significance was assessed by the Mann-Whitney Wilcoxon rank-sum test, and the Wilcoxon test was used for the comparison between the two sets of fold changes. Two-tailed *p* values were computed in all cases. *-*p*<0.05; **-*p*<0.01; ***-*p*<0.001; ****-*p*<0.0001. The list and data of all genes can be found in Tables S16 (complete data) and S18 (box plot data). **(d-e)** *unc-54*p::Q40::YFP nematodes were treated with EV or *lin-42* RNAi in plates containing either 0.1% DMSO (vehicle) or Harmine (Har, 40 μM and 60 μM). Paralysis was scored daily. n=140 worms per condition, graph is representative of three independent experiments. Significance was measured by the Log-rank Mantel-Cox test, two-tailed *p* values were computed. ****-*p*<0.0001.

**Fig. S15.**
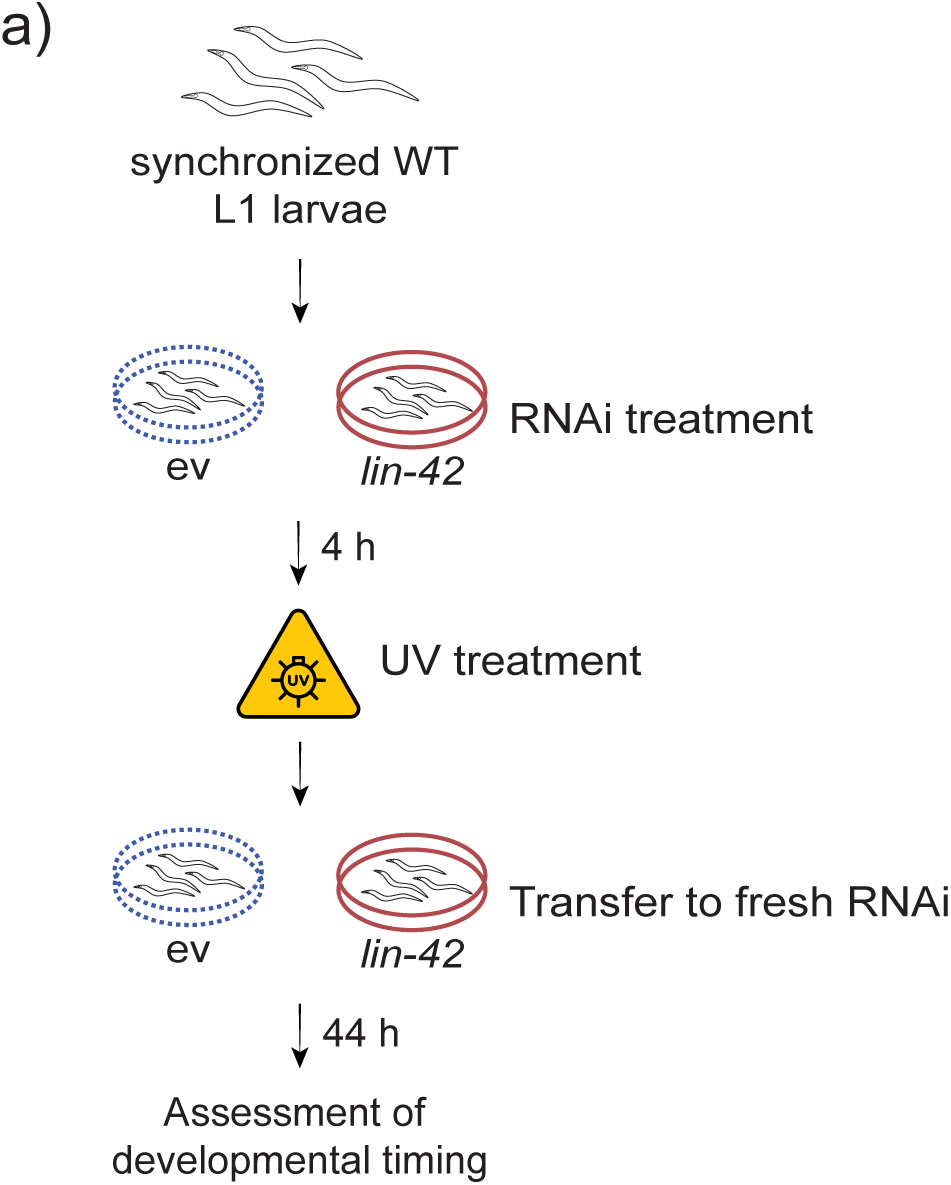
Experimental design for assessing developmental timing upon UV-B and *lin-42* RNAi treatment. **(a)** Experimental design corresponding to Fig. 6c. Age-synchronized wild-type nematodes were treated with EV or *lin-42* RNAi from L1 stage. Four hours post seeding, animals were treated with UV-B 400 or 500 mJ/ cm^2^. After treatment, worms were transferred to fresh EV or *lin-42* RNAi plates to ensure continued RNAi. Forty-four hours post UV treatment, the different larval stages were counted as L1-L2, L3, or L4 stages.

## Materials and methods

### C. elegans strains

The following *C. elegans* strains were used in this study: N2 (*C. elegans* Bristol wild-type isolate), GMC101 *dvls100 [unc-54p::Aβ_1-42_::unc-54 3’UTR+ mtl-2p::GFP]*, AM141 *rmls133 [unc-54p::Q40::YFP]*, MT2257 *lin-42(n1089)* II, RB1843 *lin-42(ok2385) II*, MRS 387 *sevIs1* [Psur-5::luc+::gfp], MOL459 *lin-42(n1089)* II; sevIs1[Psur-5::luc+::gfp], MOL460 (*lin-42(ok2385)* II; *sevIs1*[Psur-5::luc+::gfp]) YZ. GMC101, AM141, MT2257 and RB1843 strains were obtained from the Caenorhabditis Genetics Center (CGC, University of Minnesota). Luminometry strains were generated as described in^1^ and crossed to MT2257 and RB1843 strains. All strains were maintained under standard laboratory conditions as described in^2^.

### Bacterial strains and experimental conditions

*E. coli* OP50 and *E. coli* HT115(DE3) were seeded on lysogeny broth (LB) agar plates and kept at 4°C. OP50 cultures were grown overnight in LB medium without antibiotics and were used on standard nematode growth medium (NGM) agar for maintenance of all *C. elegans* strains. For RNAi experiments, HT115 bacteria expressing the specific dsRNA construct were obtained from ORF and Ahringer libraries (Source Biosciences, Nottingham, UK). All RNAi bacteria were grown in LB agar plates supplemented with ampicillin (100 mg/mL) and tetracycline (12.5 mg/mL) (Roth, #K029.1 and #0237.1). Plates were kept at 4°C, and overnight cultures were grown in an LB medium containing ampicillin. RNAi expression was induced by adding 2 mM isopropylthiogalactoside (IPTG, Sigma, #16758) and incubating the cultures at 37°C for 25 min before seeding bacteria on NGM agar supplemented with ampicillin and 3 mM IPTG. For double RNAi experiments, the OD of each culture was measured at 590 nm (Tecan, Männedorf, Switzerland) and mixed in a 1:1 ratio. Harmine (Sigma-Aldrich, SMB00461) was dissolved in DMSO and given in NGM agar in combination with ampicillin and IPTG.

### Bacterial sequencing

Plasmid DNA of RNAi bacteria was isolated using an innuPREP plasmid mini kit (Analytik Jena, Jena, Germany) and following the manufacturer’s instructions. pDNA concentration was measured by Nanodrop (Thermo Fisher Scientific, Bremen, Germany). 50-100 ng/ µL pDNA was sent together with forward and reverse primers at 10 mM (L4440 forward: 5’-GTTTTCCCAGTCACGACGTT-3’, L4440 reverse: 5’-TGGATAACCGTATTACCGCC-3’) to Eurofins Genomics (Ebersberg, Germany). RNAi hit location was verified by the Basic Local Alignment Search Tool (BLAST, Wormbase).

### Luminometry assay

We measured the duration of intermolts and molts by using a luminescence-based method^3^. To obtain synchronized eggs, 10-12 gravid hermaphrodites were transferred to fresh NGM plates to lay eggs for 1 hour. Using an eyelash, embryos were individually transferred to wells of a 96-well plate, each well containing 100 μl of S-basal (10 μg/ml cholesterol) with 200 μM Luciferin, 2 mM IPTG and 200 μg/ml Ampicillin. As food source, we added 100 μl of S-Basal containing 20 g/l bacteria containing the pL4440 empty vector for the control *(ctrl), lin-42 (n1089)* and *lin-42 (ok2385)* conditions, or with bacteria expressing RNAi against *lin-42* (*lin-42 RNAi*). To obtain the 20 g/l bacterial food source, the bacterial strains were cultured in LB medium containing 100 mg/ml Ampicilin and 12.5 mg/ml Tetracycline o/n at 37 °C, in constant shaking. The next day, the cultures were diluted 1:10 in fresh LB (with Ampicilin and Tetracycline) and incubated for 3 hours in the same conditions. After that, 1 mM IPTG was added and cultured for another 2 hours. To finally obtain the 20 g/l concentration, the bacterial cultures were centrifugated and resuspended in S-Basal twice, then adjusted the bacteria concentration was adjusted to 20 g/l. After addition of the bacteria, the plates were sealed using a gas-permeable membrane. The plate was introduced in the luminometry reader (Berthold Centro XS3), placed inside a cooled incubator (Panasonic MIR-254) set at 16.5 °C, to allow a temperature on the plate of 20 °C. We measured luminesce for 1 sec at 5-min intervals, until animals reached adulthood. We analysed luminometry data as previously described^3^.

### Lifespan analysis

N2 worms were cultivated in RNAi plates and fed with bacteria expressing RNAi against *lin-42* or harboring the empty vector (EV) control construct. Specific RNAi exposure was initiated at two different time points: i) from the L1 stage or ii) from the L4 stage. For group ii, the worms were grown in EV plates from L1 stage and until the start of *lin-42* RNAi exposure. For each lifespan condition, two plates of 70 worms were used. The viability was accessed daily by using SteREO Discovery.V8 microscope (Carl Zeiss Microscopy, Jena, Germany), and the worms were transferred to new freshly seeded plates every 2 days until adulthood day 10 (AD10), and every 4 days thereafter. Every worm that showed no touch response was considered dead. Internal hatching, gut explosions and escaping worms were censored.

### GMC101 paralysis tests

Paralysis was evaluated as previously reported^4^. *lin-42* RNAi exposure was initiated at two time points i) L1 stage or ii) L4 stage. For group ii, the animals were grown on EV expressing bacteria from L1 and until the L4 stage. Two plates of 70 worms were used for each condition. All plates were kept at 20°C until the L4 stage, and afterwards half was shifted to 25°C to trigger Aβ toxicity. Paralysis was scored daily using SteREO Discovery.V8 microscope (Carl Zeiss Microscopy, Jena, Germany). Immobile worms insensitive to touch were considered paralyzed. Worms were transferred to new freshly seeded plates every second day until AD10, and every fourth day thereafter.

### AM141 paralysis tests

Paralysis was evaluated as previously reported^5^. *lin-42* RNAi exposure was initiated at two time points i) L1 stage, ii) L4 stage. For group ii, the animals were grown on EV expressing bacteria from L1 and until the L4 stage. Two plates of 70 worms were analyzed for each condition. Paralysis was observed using SteREO Discovery.V8 microscope (Carl Zeiss Microscopy, Jena, Germany). Immobile worms insensitive to touch were considered paralyzed. Worms were transferred to new freshly seeded plates every second day until AD10, and every fourth day thereafter.

### Proteomics sample collection, data acquisition and analysis

Age-synchronized populations of N2 (wild type) worms were exposed to specific RNAis and their combinations i) from the L1 stage and ii) from the L4 stage and collected at adulthood day 2 (AD2); 800 worms per condition were collected. For group ii, worms were grown on EV RNAi from L1 stage and until L4 stage. Sample, preparation, data acquisition and data analysis (DIA mode) were performed as described in detail previously^2^. The data (candidate table) and data reports (protein quantities) were exported, and further data analyses and visualization were performed with Rstudio using in-house pipelines and scripts.

### Fluorescence microscopy

AM141 (Q40-YFP) animals treated as described in the figures were imaged at L4, AD1, and AD2 stages using Axio Zoom.V16 microscope fitted with PlanApo Z 0.5x objective (Carl Zeiss Microscopy, Jena, Germany). Fixed exposure times of 150 ms and 4.8 ms were used for YFP and brightfield imaging respectively, and 90x magnification was applied. Worms were immobilized by placement on ice for 30 min prior to microscopy, and 2 plates of 15 animals were scored for each condition. Image analysis was carried out using Zen software v2.6, blue edition (Carl Zeiss Microscopy, Jena, Germany). The area (body size, head to tail), YFP intensity and number of aggregates were measured for each animal, and following calculations were made: YFP intensity/area and number of aggregates/YFP intensity.

### Western blotting

800 AD2 stage N2 worms per condition were harvested in 1xM9, followed by extraction with the lysis buffer (100 mM Hepes pH 8.5, 100 mM DTT, 1 mM EDTA, 1% SDS, 1 mM PMSF, 1x Protease inhibitor cocktail (Roche, #04693116001)) and homogenized by using Precellys 24 device and tubes (Precellys, Bertin Technologies, France). Four homogenization cycles (20 sec at 6.000 rpm) were applied separated by a 30-sec interval. The samples were centrifuged at 2500 rpm for 30 sec into a 1.5 mL tube, then sonicated (Bioruptor Plus, Diagenode, Belgium) for 10 cycles (60 sec ON/ 30 sec OFF) at a high setting and at 20°C. After sonication, samples were boiled at 95°C for 5 min and centrifuged at 10600 rpm for 10 min.

Protein content was quantified using the EZQ Protein Quantitation Kit (Molecular Probes, Thermo Fisher Scientific, #R33200). Then, 20 ug protein was separated in 4-20% Mini-Protean TGX Precast Gels (Bio-Rad, California, US, #4561094) and blotted using Bio-Rad Trans-Blot Turbo device (Bio-Rad, California, US) onto a Nitrocellulose membrane 0.2 µm (Bio-Rad, California, US) for 30 min. Proteins were visualized with Ponceau S staining before incubation with 5% BSA (AppliChem, Darmstadt, Germany) in TBST for 1h at room temperature. Membranes were then incubated with primary antibodies in 5% BSA in TBST overnight at 4°C. The following primary antibodies were used: total histone H1 (Invitrogen, #PA5-30055, rabbit, 1:1000), total histone H2AZ (Abcam, #ab150402, rabbit, 1:1000), total histone H2B1L (St John’s Laboratory, #STJ195934, rabbit, 1:1000), total histone H3 (Sigma, #H0164, rabbit, 1:1000), total histone H4 (Merck, #07-108, rabbit, 1:1000), alpha-tubulin (Sigma-Aldrich, #T6199, mouse, 1:1000). After 12h, the membranes were washed 3 times with TBST for 5 min and incubated at room temperature with a secondary antibody: IRDye anti-mouse 1:10 000 (LI-COR, Lincoln, US, P/N 926-322210 and P/N 926-68070) and IRDye anti-rabbit (LI-COR, Lincoln, US, P/N 926-32211 and P/N 926-68071) 1:10 000 for 1h. Following this step, membranes were washed 3 times with TBST for 5 min and 2 times with TBS for 10 min. Detection was carried out using an Odyssey device (LI-COR, Lincoln, US) and quantification was performed using Image Studio Lite (Version 5.2.5). All histone signals were normalized by using corresponding alpha-tubulin signals.

### RT-qPCR with nematode samples

250 synchronized AD2 stage N2 animals were harvest in 1x M9 followed by resuspension in 1 mL Tri Reagent (Sigma Aldrich, Darmstadt, Germany). Samples were homogenized using Precellys 24 device (Precellys, Bertin Technologies, France) with 2 cycles of 20 seconds/6.000 rpm, separated by a 30-sec interval. Subsequently, 5 min incubation at room temperature with 100 µL 1-Bromo-3-chloropropane (BCP) (Merck, Darmstadt, Germany, #B9673) was performed. Samples were vortexed for 15 sec and incubated for 2 min at room temperature. Then, centrifuged at 12.000 rpm for 15 min at 4°C. The upper clear phase was collected and mixed with an equal amount of 70% EtOH in RNAse-free water (HyPure, Cytiva, Utah, US). RNA isolation was performed by using an innuPREP RNA mini kit (Analytik Jena, Jena, Germany) and following the manufacturer’s instructions. The samples were further treated with DNase and RNase (Thermo Fisher Scientific, Bremen, Germany) by following the manufacturer’s instructions for a reaction with 1 µg total RNA. Reverse transcription was done with 0.5 µg total RNA using RT SuperScript III kit (Invitrogen, California, US) and following the manufacturer’s instructions. RT-qPCR reactions were performed in triplicates in 96-well plates, using SSoAdvanced SYBR Green master mix (Bio-Rad, California, US) and following the manufacturer’s instructions in a Bio-Rad CFX96 touch real-time machine (Bio-Rad, California, US). The relative gene expression was calculated by using the 2^−ΔΔCt^ method (Livak & Schmittgen, 2001). The list of primers can be found in Table S24.

### Analysis of DREAM binding sites at candidate genomic regions

The previously published ChiP-Seq dataset was obtained from^6^, and overlapping genome binding sites of the three DREAM subunits mediating the health effects of circadian dysfunction (LIN-9, LIN-53, and LIN-54) were searched against promoter regions of candidate genes by using the *C. elegans* genome assembly version WS220/ce10 (UCSC Genome Browser Gateway database).

### Cell culture, siRNA exposure and Harmine treatment

RPE-1 hTERT cells (#RPE CRL-4000, ATCC) were cultured in DMEM media with F12 (#31331093, Thermo Fisher Scientific). Culture media were supplemented with 10% fetal bovine serum (FBS; # A5209402 Thermo Fisher Scientific) and penicillin/streptomycin (#15140122, Thermo Fisher Scientific). Cell lines were tested at least twice a year for *Mycoplasma* contamination using the LookOut Detection Kit (Sigma), and all tests were negative.

Cells were treated with DMSO (0.1%; Carl Roth, Karlsruhe, Germany) or Harmine (10 μM; Sigma-Aldrich, SMB00461) for 20 h. For knockdown experiments, cells were reverse transfected with 10 nM Silencer Select siRNAs (Thermo Fisher Scientific) using RNAiMAX (#13778150, Fisher Scientific) and Opti-MEM (#31985062, Thermo Fisher Scientific) following the manufacturer protocol. The following siRNAs were used (Thermo Fisher Scientific): siControl (#4390844) and siPER1 (#s10301).

### Reverse transcription semi-quantitative real-time PCR (RT-qPCR) in RPE cells

Total cellular RNA was extracted using the innuPREP RNA Mini Kit (Analytik Jena, Jena, Germany) following the manufacturer protocol. One-step reverse transcription and real-time PCR was performed with a Quantstudio 5 using Power SYBR Green RNA-to-CT 1-Step Kit (Thermo Fisher Scientific) following the manufacturer protocol. The list of primers used can be found in Table S21.

### RNA-sequencing in RPE cells

Total RNA was extracted from 5 replicas using the innuPREP RNA Mini Kit (Analytik Jena, Jena, Germany) following the manufacturer protocol. Sequencing of RNA samples was performed using Illumina’s next-generation sequencing methodology^7^. In detail, total RNA was quantified, and quality checked using Tapestation 4200 instrument in combination with RNA ScreenTape (both Agilent Technologies). Libraries were prepared using a Biomek i7 by introducing 300 ng of total RNA using NEBNext Ultra II Directional RNA Library Preparation Kit in combination with NEBNext Poly(A) mRNA Magnetic Isolation Module and NEBNext Multiplex Oligos for Illumina (Unique Dual Index UMI Adaptors RNA) following the manufacturer’s instructions (New England Biolabs). Quantification and quality check of libraries was done using a Tapestation 4200 instrument and a D1000 ScreenTape (Agilent Technologies). Libraries were pooled and sequenced on a NovaSeq6000 SP 100 cycle run. System ran in a 101 cycle/single-end/standard loading workflow mode. Sequence information was converted to FASTQ format using bcl2fastq v2.20.0.422.

Raw read quality trimming (5nt sliding window with cutoff 20) was performed using Trimmomatic v0.39^8^ and TruSeq universal adapter were clipped (minimum overlap 5nt) with Cutadapt v2.10 (minimum length 18)^9^. Potential sequencing errors were detected and corrected utilizing Rcorrector v1.0.4^10^ prior to artificial rRNA depletion using SortMeRNA v2.1b^11^. Reads were aligned to the human genome with segemehl v0.3.4^12, 13^ (split-read mode, accuracy 95%, reference GRCh38) and filtered for unique alignments with SAMtools v1.14^14^.

Alignments have been deduplicated for over amplified PCR fragments based on unique molecular identifiers utilizing UMI-tools v1.1.1^15^ and subsequently quantified with featureCounts v2.0.1 (10nt minimum exon based meta-feature overlap, reference annotation Ensembl v106)^16^. Differences (log2FC) between conditions mentioned in the figures were calculated with DESeq2 v1.34.0^17^.

### Melatonin treatment of mice and organ collection

Application UKJ-18-036 “CLOCK”. 10-week-old C57BL/6 mice were housed under a 12h/12h light cycle, with controlled temperature. Melatonin (100 mg/kg) was administered via drinking water as described previously (Rajput et al., 2017) for 2 weeks. Melatonin was first diluted in ethanol and the final concentration was achieved by diluting with drinking water (e.g. 8 mg melatonin in 100 µL alcohol, final alcohol concentration is less than 0.5%). The same amount of ethanol in the drinking water without melatonin was used as a control (vehicle). After melatonin administration tissue (hippocampi) was collected, with half of the animals sacrificed in the active phase (dark phase, night) and half in the inactive phase (light phase, day).

### Gene expression analysis of mouse brain tissue by qPCR

RNA was isolated using Trizol (#15596026, Thermo Fisher) resuspension followed by chloroform precipitation. The RNA was then washed with isopropanol and 75% ethanol. cDNA synthesis was carried out using the GoScript™ Reverse Transcriptase kit (#A5001, Promega). Quantitative real-time PCR was subsequently performed on a qTower3GTouch machine from Analytik Jena using SsoFast EvaGreen Supermix (#1725201, Bio-Rad). Details of the primer sequences can be found in the supplementary material (Table S22). The quantitative PCR data were analyzed as previously published (Trautmann C et al, 2020).

### Mouse sleep deprivation, tissue collection and RNAseq (reanalysis Jan *et al.*, 2024)

As described in Jan *et al.*, 2024 sleep deprivation (SD) was performed by gentle handling (Mang & Franken, 2012) for 6h starting at light onset (zeitgeber time ZT0-6). Mice were anesthetized with isoflurane prior to decapitation. Cortex and liver were rapidly dissected, and flash frozen in liquid nitrogen. Time schedule of tissue sampling was described (Hor et al., 2019). Frozen cortex samples were processed as described (Hor et al., 2019). RNAseq cortex data from Jan *et al.*, 2024 were obtained through the GSE262410 and the expression of DREAM subunits and DNA damage and repair genes were filtered. The log_2_ CPM values were then used to visualize sleep versus wake expression dynamics. The complete RNAseq data from data points NSD ZT3 and SD ZT27 (ZT3) and NSD (ZT6) and SD ZT30 (ZT6) were used to make the relative difference between sleep and sleep deprivation. The log2 values were used for further analyses and visualization with Rstudio using in-house pipelines and scripts.

### Statistical analysis

The raw data of all experiments were plotted in Excel sheets, GraphPad Prism, or Rstudio files. The tests used in each experiment are mentioned in respective figure legends. The differences were considered significant with *p < 0.05, **p < 0.01, ***p < 0.001, n.s. = not significant, unless otherwise stated.

